# What drives intersubject correlation of EEG during auditory narratives?

**DOI:** 10.64898/2026.02.19.706583

**Authors:** Emilia Fló, Álvaro Cabana, Juan Valle-Lisboa, Damian Cruse, Jens Madsen, Lucas C. Parra, Jacobo Sitt

## Abstract

When participants are engaged with auditory narratives, physiological and neural signals exhibit temporal correlations between subjects. The intersubject correlation (ISC) increases when attention is directed to the stories, suggesting that shared neural and bodily dynamics arise from a similar processing of the narratives. In this study, we investigated whether the ISC of the EEG elicited by auditory narratives is driven by low-level acoustic (envelope, spectrogram) and/or higher-level linguistic information (word onset, word surprisal) in two groups of healthy participants during passive, attentive and distracted listening. We use temporal response functions (TRFs) for acoustic, and linguistic features to assess the contribution of each feature to the ISC, measured using correlated component analysis (CorrCA). TRFs derived for acoustic features explained a larger fraction of variance in the EEG than linguistic features and were the main contributors to the ISC. The attention-related increase in ISC was driven by all features. Importantly, word surprisal had an effect on ISC only during active story engagement, with timing and scalp distribution consistent with language processing. Notably, the linear responses captured by TRFs only explained a small amount of the overall ISC, suggesting that ISC is largely driven by nonlinear responses to the narratives. Identifying the factors that drive these common neural responses, is clinically relevant for interpreting EEG ISC exhibited in unresponsive patients. We propose that the combined use of ISC and TRFs has the potential to provide meaningful markers of language processing in patients with disorders of consciousness, and we suggest practical recommendations for implementing these approaches in clinical settings.

## Introduction

When participants are exposed to auditory narratives, heart rate, pupil size, and neural activity show a temporal correlation between subjects (Hasson et al., 2004; Madsen & Parra, 2022; Naci et al., 2014; Pérez et al., 2021; Türker et al., 2023; Wilson et al., 2008). In EEG recordings, this intersubject correlation (ISC) is increased when attention is directed to the stories (Ki et al., 2016; Madsen & Parra, 2022, 2024; Rosenkranz et al., 2021), suggesting that the source of covariation in brain and bodily signals arises from a similar processing of narratives. The synchronization of brain activity in EEG can be measured using correlated component analysis (CorrCA), a statistical procedure that reduces the dimensions of EEG data from electrodes to a subset of components that maximize the correlation of the evoked responses across all subject pairs (Parra et al., 2019). It has been suggested that the resulting ISC components of the EEG reflect sensory processing (Cohen & Parra, 2016; Poulsen et al., 2017), are modulated by attention (Ki et al., 2016), capture varying levels of engagement (Cohen and Parra 2016; Dmochowski et al. 2012; Poulsen et al. 2017), and predict recall of the narrative (Cohen and Parra 2016). Nevertheless, to our knowledge, no study has examined the contribution of specific features of the speech stimuli to the ISC, nor how attention modulates their effects.

Speech is a complex natural stimulus for which humans are specially tuned to parse and comprehend. The brain processes involved in understanding speech are likely shared among individuals and could be driving the observed common neural responses to auditory narratives. Speech is a continuous signal with a hierarchical structure such that sentences can be decomposed into words, words into phonemes, and phonemes into moment-to-moment spectral variations, and this hierarchy is reflected in the spatial and temporal unfolding of speech processing in the brain (de Heer et al., 2017; Hickok & Poeppel, 2007; Lerner et al., 2011). Speech comprehension thus requires the brain to decode and integrate information across these multiple levels of information as they unfold continuously over time.

Linear encoding models (Holdgraf et al. 2017) can be used to study the relationship between multiple embedded levels of information in complex naturalistic stimuli and neural activity. In our case, properties of the speech signal can be extracted from the stimulus to train a model on a given feature or set of features. Then, a stimulus-response function is obtained that can be used to predict the neural response for a new stimulus. Here we use temporal response functions (TRFs) (Crosse et al., 2016; Ding et al., 2012), linear encoding models that take into account the delays in neural processing. TRFs quantify the linear dependency between neural activity and a speech feature, which can be interpreted as a measure of how well the evoked neural response is time-locked to the speech feature (Brodbeck & Simon, 2020). This method demonstrates a temporal locking of EEG to the acoustic features of speech such as the envelope (Prinsloo & Lalor, 2022; Rosenkranz et al., 2021), the spectrogram (Di Liberto et al., 2015; Ding et al., 2012; Teoh et al., 2022), the spectral information of individual phonemes (Daube et al. 2019), and phoneme categories (Di Liberto et al., 2015; Di Liberto & Lalor, 2017; Teoh et al., 2022). In addition, TRFs are sensitive to linguistic features such as the semantic dissimilarity of a word relative to the previous context (Broderick et al., 2018, 2019, 2022), word and phoneme segmentation (Gillis et al., 2021, 2023; Teoh et al., 2022), and lexical properties of words (Gillis et al., 2021, 2023). Directing attention to speech enhances the responses to these features, suggesting a stronger cortical tracking of the features when participants are engaged with the stimuli (Broderick et al., 2018; O’Sullivan et al., 2015; Power et al., 2012; Teoh et al., 2022). Importantly, EEG evoked responses to linguistic features show a stronger correlation to speech comprehension than responses to acoustic information (Shyanthony R. Synigal et al., 2023).

In this study, we use univariate and multivariate TRFs (mTRFs) to investigate the contributions of low-level acoustic information and higher linguistic features to the intersubject correlation elicited during passive listening without task-related demands and during attentive and distracted listening to auditory narratives. Finally, we discuss the potential of combining these tools to assess language processing and awareness in non-communicative patients and provide some methodological suggestions for a successful implementation.

## Materials and Methods

### Experiment 1: passive listening to auditory narratives

#### Participants

Twenty-seven native English speakers (22 females, mean age = 21.20, sd = 1.85, range= 18-26) participated in this study. The experiment was conducted at the Centre for Human Brain Health, University of Birmingham, England, and was approved by the STEM ethics committee of the University of Birmingham, England. All subjects provided written informed consent.

#### Stimuli and procedure

The participants listened to a 16-minute extract of the audiobook 20,000 Leagues Under the Sea (author: Jules Verne, read by: David Linski, male speaker, British English accent, public Domain (P) 2017 Blackstone Audio, Inc.). The audiobook extract was taken from the first chapter and half of the second chapter. The story was presented in segments of 1 minute each, yielding 16 trials sampled at 16 kHz and presented at a comfortable listening level, which was individually adjusted for each participant. The instructions given to the subject were ‘to listen to the story and look at a fixation cross’. The stimuli were delivered by headphones - ER1 Insert Earphones (Etymotic Research), using Psychopy v3.1.2 (Peirce, 2007).

#### Recording and preprocessing

EEG was recorded with 128 channels with a Brainvision amplifier with a sampling frequency of 250 Hz referenced to CPz. Heart activity was also recorded, and results have already been published (Pérez et al., 2021). Data was filtered between 0.1 and 40 Hz with a bandpass filter (one-pass zero-phase FIR filter with a length of 8251 samples). Channels were rejected if their variance was above 3.5 standard deviations from the mean channel variance, this was iteratively performed 4 times. ICA was performed to remove ocular artifacts, bad channels were interpolated, and data were segmented into 16 trials. Sparse artifact removal (De Cheveigné, 2016) was carried out using the meegkit library (https://github.com/nbara/python-meegkit). Finally, EEG channels were combined into a 64-channel Biosemi layout configuration to be comparable to Experiment 2 and referenced to the average of the 64 channels. For EEG preprocessing MNE 1.0.3 (Gramfort, 2013) was used.

### Experiment 2: attentive and distracted listening of auditory narratives

#### Participants

Thirty-two English speakers took part in the experiment (16 females, mean age = 23.69, sd = 4.42, range = 19-36, 3 subjects were removed due to bad signal quality or issues during stimulus presentation, and 3 other subjects were removed for not completing the task). Experiments were carried out at the City College of New York with the approval of the Institutional Review Boards of the City University of New York. All subjects provided written informed consent before the experiment.

#### Stimuli and procedure

The stories consisted of 10 narratives selected from the StoryCorps project aired on National Public Radio (NPR): “Eyes on the Stars” (one male speaker, American English accent), “John and Joe” (one male speaker, American English accent), “Marking the Distance” (one male and one female speaker, American English accent), “Sundays at Rocco’s” (one male speaker, American English accent,) and “To R.P. Salazar with Love” (one male and one female speaker, American English accent) and the New York Times Modern Love series: “Broken Heart Doctor” (one male speaker, American English accent), “Don’t Let it Snow” (one male and one female speaker, American English accent), “Falling in Love at 71” (one female speaker, American English accent), “Lost and Found” (two female speakers, American English accent), and “The Matchmaker” (one male speaker, American English accent). Each story was 2–5 min in duration and was presented as continuous narration sampled at 48 kHz and presented at a comfortable listening level, adjusted for each participant The total duration of the auditory stimuli was 27 min. The stories were played through stereo speakers placed at 60° angles from the subject while facing a gray background in a 27” monitor placed at a distance of 60 cm from the participant. In the attentive condition, participants were asked to attend to the story while looking at a fixation cross. Comprehension questions were asked after each story, and all participants achieved more than 50% accuracy overall (mean accuracy = 77.88%, sd = 11.06%, range = 51.39–98.61%). In the distracted condition, participants listened to the stories again, but they were asked to count backwards silently in their mind in steps of 7 starting from a random prime number between 800 and 1000 (distracted condition). All participants listened to the stories first in the attentive condition, followed by the distracted condition.

#### Recording and preprocessing

Detailed preprocessing for this data set can be found in (Madsen & Parra, 2024). Briefly, EEG was recorded with 64 channels in a 10/20 configuration at a sampling frequency of 2048 Hz using a BioSemi Active Two system. In addition, the electrooculogram (EOG) was recorded with external electrodes placed at the left and right external ocular canti, and under and above the right eye. The signals were digitally high-pass filtered (0.01 Hz cutoff) and notch-filtered at 60 Hz to remove line noise. Signals were digitally low-pass filtered (64 Hz cutoff) and downsampled to 128 Hz. Bad electrodes were identified manually and replaced with interpolated channels. The EOG channels were used to remove eye-movement artifacts. Finally, data was epoched into the 10 stories and a 0.1:40 Hz bandpass filter was applied before referencing the data to the average of the 64 channels.

### Intersubject correlation

Correlated component analysis (CorrCA) (Cohen & Parra, 2016; Dmochowski et al., 2012; Parra et al., 2019) extracts projections of the data with maximal correlation across repetitions, in our case maximal correlation of EEG-evoked response between participants listening to the same audio. CorrCA yields a weighted combination of channels that maximally separate the between subject covariance (*R_b_*) from the within subject covariance (*R_w_*) across trials **(Fig. 1A).** For Experiment 1 the number of trials was 16, for Experiment 2 each story was split into two segments yielding 20 ‘trials’ (this was carried to increase the number of folds for TRF fitting, see below). Matrices *R_w_* and *R_b_* are averaged over trials (all trials for Experiment 1, and all trials including both conditions for Experiment 2), and the eigenvectors (*V*) of the matrix *R_w_* ^−1^*R_b_* are calculated as follows:

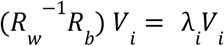

**Fig 1.**
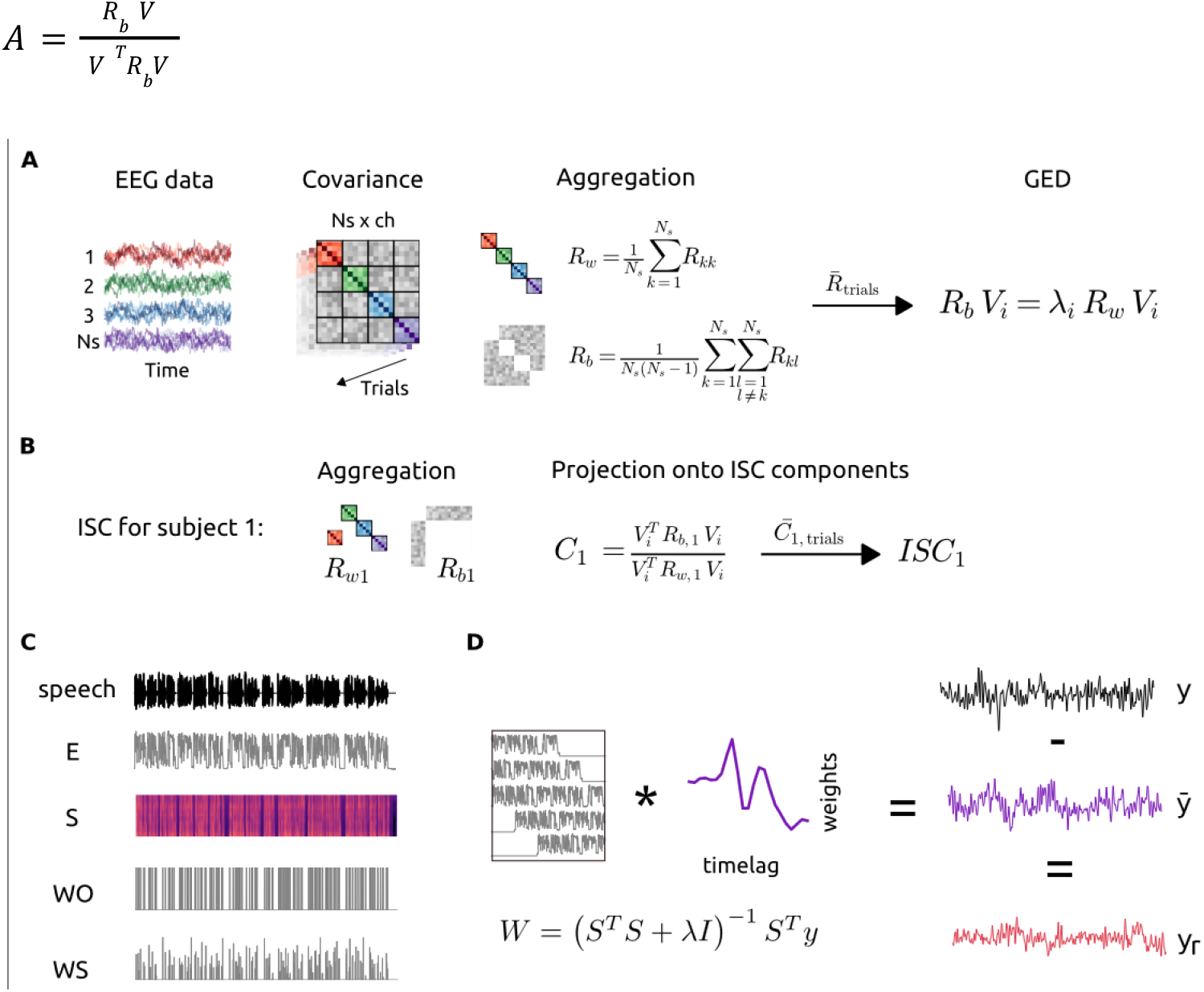
**A**. Correlated component analysis. For each trial, cross-covariance is computed for each pair of electrodes and each pair of subjects (i, j, with i ≠ j and i, j ∈ {1, …, N◻}), and aggregated resulting in a between-subject covariance matrix (*R_b_*) and a within-subject covariance matrix (*R_w_*). These matrices are averaged across trials, and the component vectors (*V_i_*) of the matrix *R_w_* ^−1^ *R_b_* are obtained using generalized eigenvalue decomposition (GED). **B.** To illustrate the procedure, we describe the computation of subject-level intersubject correlation (ISC) for Subject 1. ISC is obtained by averaging the between-subject covariance and the within-subject covariance, for a given stimulus and condition, across all subject pairs that include Subject 1. Pearson correlation between the evoked responses of Subject 1 and those of the remaining participants (*C*_1_) is computed by projecting these covariance matrices onto the component vectors *V*. Intersubject correlation for subject 1 (*ISC*_1_) is then obtained by averaging correlation values across trials within the same experimental condition (*C*_1,*trials*_). **C**. Representation of the features extracted from the speech signal. E - envelope, S - spectrogram, WO - word onset, WS - word surprisal. **D**. Temporal response functions (TRFs) were estimated using regularized linear regression from time-lagged copies of each feature (S), as represented by the envelope in this example. The TRFs predicted responses (*y*) were obtained for each trial and feature. The predictions were subtracted from the actual EEG response and the residual activity (*y_r_*) was obtained. The ISC was computed for the original data and for the residual activity.

The eigenvectors *V_i_* with the strongest eigenvalues λ*_i_* define the dimensions that capture the largest correlation between subjects. For both datasets, the covariance matrix *R_w_* was regularized using a shrinkage parameter (γ = 0.5). Pearson correlation of evoked responses for a given participant to the rest of the group experiencing the same stimulus in the same condition was obtained by projecting the data onto the vectors *V_i_*

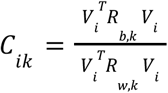

where *R_b_*_,*k*_ represents the between-subject covariance and *R_w_*_,*k*_ the within-subject covariance for a given stimulus and condition, averaged across all pairs of subjects involving a given subject. The individual intersubject correlation for each component and condition was obtained by averaging the correlation values for all trials during the same experimental condition **(Fig. 1B)**. Here, we only report the correlation values for the first three components that show values significantly different from zero (for component three the correlations for most participants are not statistically different from chance) (**Fig. S2**). The ISC was computed using Matlab R2021a and modifying custom scripts available at http://www.parralab.org/isc/. For a detailed description of the method see (Parra et al. 2019). Finally, to see the contributions of each electrode to the components we compute a forward model (Haufe et al., 2014) as:

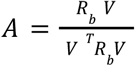

### Speech features extraction

The broadband envelope (E) was obtained for each speech stimulus using the function mTRFenvelope of the mTRF-Toolbox which estimates the root-mean-square envelope by squaring the signal, low-pass filtering it over a sliding window, and taking the square root. To model human hearing, the output was compressed by raising the value to 0.3 (Biesmans et al. 2017; Crosse et al. 2021). The envelope was computed for a 50 Hz sample rate for Experiment 1 and 64 Hz for Experiment 2. The spectrogram (S) was obtained by filtering the stimuli into 16 bands between 250 Hz and 8 kHz, and computing the envelope for each band as the absolute value of the complex analytical signal given by the sum of the original signal and its Hilbert transform, resulting in a logarithmically scaled Mel-spectrogram (Schädler & Kollmeier, 2015). Signals were resampled following the envelope computation. To obtain word onsets, audio files were transcribed using Whisper (https://openai.com/research/whisper), manually corrected, and forced aligned to the speech signals using Montreal Forced Aligner 2.0 (McAuliffe et al., 2017). Word onset (WO) was coded as a binary array with value one whenever the onset of a word occurred. The BERT language model pre-trained with RoBERTa base corpus (Liu et al., 2019) was used to obtain word predictability values using a strictly preceding-context procedure. Specifically, for each target word, the model was provided only with the words preceding that word, followed by a single mask token, all subsequent words were excluded from the input. For Experiment 1, the context was determined by the 420 previous words and for Experiment 2, the 350 previous words of each story were used as context. The probability assigned to the target token at the masked position was extracted from the model output. For words split into multiple subword tokens, only the probability of the first subword token was used. Word surprisal (WS) was defined as − *log*2(*p*), where *p* is the probability of the target token given by the model (**Fig. 1C**). We evaluated several autoregressive language models, including GPT-based models, but none outperformed BERT in predicting the EEG responses in our datasets. We therefore retained the original BERT-based approach for the analyses reported here. Finally, to test the validity of our WS representations in Experiment 2, a randomized WS vector was constructed (WSr) by permuting the WS values across words within stories. Segments of music at the beginning and end of stimulus in Experiment 2, and the corresponding EEG data, were not used for the analyses.

### Temporal response functions (TRFs)

A linear forward modeling approach was used to predict the EEG response given each speech representation or a combination of multiple representations using the mTRF-Toolbox (Crosse et al., 2016). Each EEG channel response was estimated as a linear convolution of the speech representations over a range of time lags relative to stimulus onset. Fitting the linear models corresponds to finding a set of weights *w* that minimize the mean squared error between the original response *y* and the one predicted by the model *y* which can be estimated using ridge regression (**Fig. 1D**). Subject-level models were computed using a leave-one-out nested cross-validation procedure, both to select the optimal regularization parameter (λ) and to evaluate prediction accuracy. For each subject, we iteratively selected one trial to test the TRF, and used the remaining trials to carry a leave-one-out cross-validation to obtain the ridge parameter across channels and folds that maximized the correlation between *y* and *y*. The optimal λ was then used to train the model on the n−1 trials, and the trained model was tested on the left-out trial (**Fig. S1**). This procedure yielded one Pearson correlation coefficient per trial and subject, computed between the predicted and the observed EEG and averaged across channels. The cross-validation was performed across regularization values in the range 10^−6^: 10^6^. The time lags used to train the model during cross-validation were −100:800 ms and the final lags for model training and testing were restricted to −50:750 ms (Crosse et al., 2021). Because the resulting TRF weights depend on the selected regularization parameter λ, we adopted a common λ value to enable comparisons across conditions in Experiment 2. Specifically, we computed the mode of the optimal λ values obtained across channels and subjects, and then recomputed the subject-level models for the WS, WO, and WSr conditions using this fixed ridge parameter. In addition, multivariate models were constructed with combinations of E, S, WO, and WS, to assess language processing beyond acoustics and word segmentation. The predicted response for each model was subtracted from the EEG and ISC was recomputed on the residual EEG.

### Statistical analyses

For each ISC component, chance level values were determined by recomputing ISC over 1000 surrogate datasets generated by circularly shifting each subject’s time series independently within each trial. Statistical significance was assessed against this null distribution and corrected for multiple comparisons across subjects using false discovery rate (FDR, Benjamini & Hochberg, 1995). Correlations between ISC and TRF accuracy across subjects were assessed through Pearson correlations. Subject-level TRF significance was evaluated using an analogous surrogate-based procedure, in which 1000 surrogate models were obtained by circularly shifting the feature–EEG alignment using the optimal ridge parameter (λ) for each subject and model. The group analyses of the TRF weights were carried out using a cluster permutation approach. Specifically, we implemented a one-way paired samples t-test using a cluster-level statistical permutation test on the TRFs for time points 0.3:0.7 s as we expected TRFs for WO and WS to show more negative values for the canonical N400 during attention to the stories. The t-test threshold corresponding to an alpha value of 0.05 was used, and samples exceeding this threshold were clustered according to temporal and spatial proximity. The t-values in each cluster (t-sum) are summed and compared to a distribution of clusters obtained from 2000 repetitions of the analysis with the condition labels randomly swapped. The proportion of clusters from the null distribution with more extreme values than the cluster obtained from the empirical data yielded the p-value for a given cluster. The level of significance was established at ɑ = 0.05.

To compare the accuracy of the models and the level of ISC between and within attentional conditions we used two-sided Wilcoxon signed-rank tests. We report the test statistic W, and the p-values yielded were FDR-corrected. Bayesian paired-samples analyses were performed. Within-subject ISC differences were tested against a null of zero mean difference using a JZS prior (r = 0.707), Evidence was quantified as BF₁₀, representing the relative support for the alternative hypothesis over the null. Statistical analyses were conducted in R (RStudio Team, 2016).

## Results

In the first experiment, we sought to determine which are the features of speech that drive the intersubject correlation observed when participants listen passively to narratives. For this, 16 minutes of an audiobook were presented to participants and encoding models for acoustic properties (envelope and spectrogram) and linguistic features (word onset and word surprisal) were obtained for each subject. The intersubject correlation was computed employing CorrCA for the original data and after removing the predicted responses for each of the encoding models. In a second experiment, we examined how attentive and distracted listening affects the ISC and the relative contributions of acoustic and linguistic features. In addition, multivariate models were constructed to account for the correlation across features.

### Intersubject correlation is elicited during passive listening of narratives

The evoked brain responses of individual participants during passive listening was similar to that of the group as measured by the ISC of the three strongest correlated components. The corresponding forward model of the components follows the distribution reported in previous studies (Cohen & Parra, 2016; Rosenkranz et al., 2021) (**Fig 2A**). ISC was above chance for all subjects for the first two components, however ISC values for the third component were very small, and after FDR correction, they did not differ from chance in half of the participants (**Fig S2**). Considering this, we focused the following analyses on the first two components.

**Fig 2.**
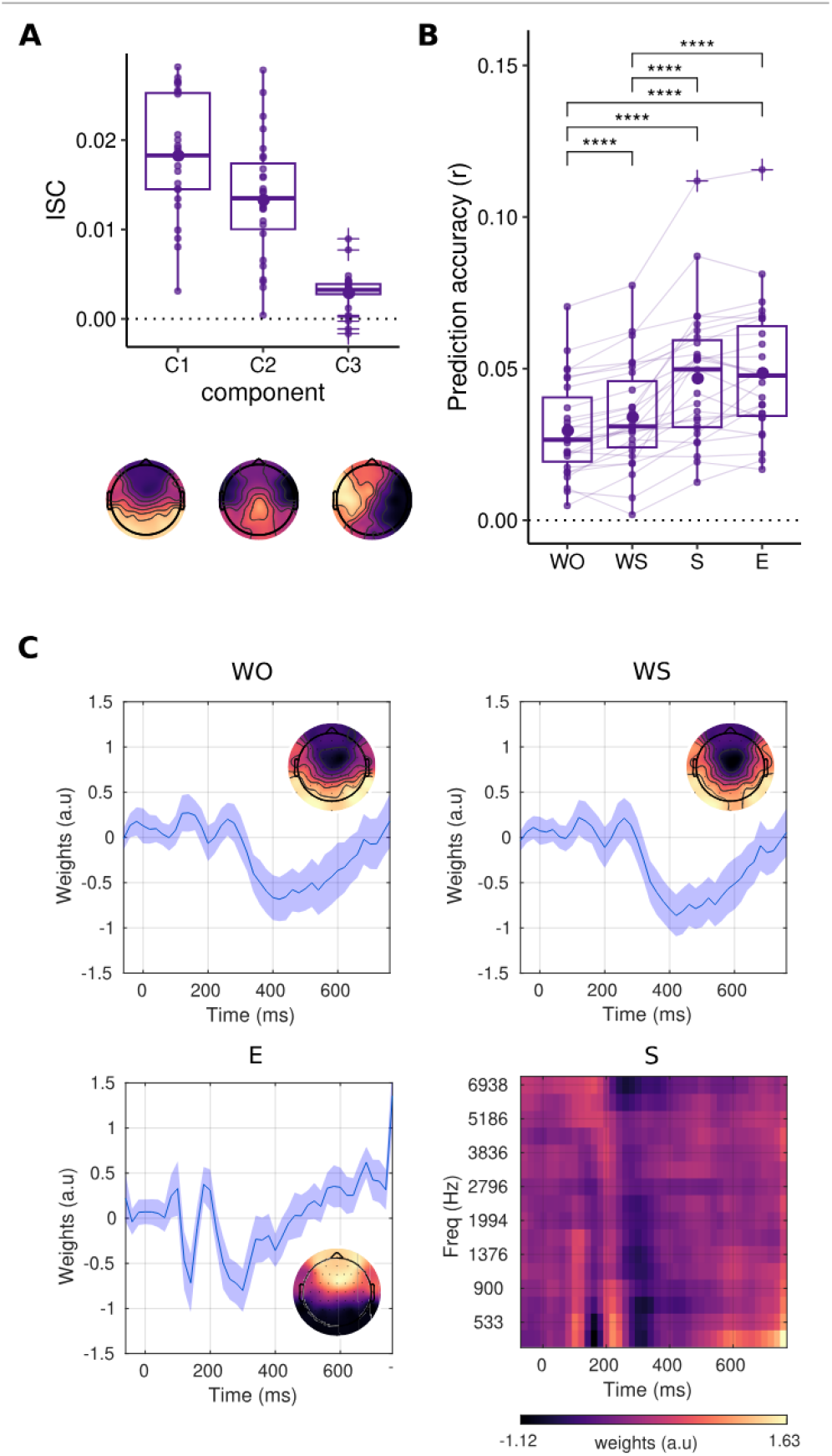
Intersubject correlation and encoding models during passive listening. **A**. Top. ISC values for the first three strongest components. Bottom. Forward model for the components capturing its correlation with each electrode. **B**. Prediction accuracy for each univariate model and subject obtained as the correlation between the predictions and the actual EEG response (WO: word onset, WS: word surprisal, E: envelope, S: spectrogram). Two-tailed Wilcoxon signed-rank test, (*) p < 0.05, (**) p < 0.01, (***) p < 0.001, and (****) p < 0.0001, FDR corrected. **C**. Univariate TRF weights normalized and averaged across subjects for Cz. The inset topographies shown correspond to the 440:480 ms time window for the WO and WS models, and 80:100 for the E model. Topographic maps show component coefficients (CorrCA) and normalized TRF weights, which represent relative spatial patterns rather than absolute physiological quantities (e.g., voltage or power).

### Both acoustic and linguistic information contribute to the ISC during passive story listening

Univariate models based on acoustic features yielded higher prediction accuracies than those based on word onset and word surprisal (mean r: E = 4.89e-2, S = 4.71e-2, WS = 3.44e-2, WO = 3.00e-2. See **Fig. 2B, Table S1** for comparisons). No differences were found in prediction accuracy between the spectrogram and envelope models (W = 138, p = 0.229), Although word surprisal and word onset features are highly similar and therefore their corresponding TRFs predicted largely overlapping EEG responses, word surprisal models were more accurate (W = 31, p = 7.04e-5), and a larger proportion of subject-level WS TRFs reached significance (**Fig. 3A**). The temporal profiles of the TRFs weights indicated that acoustic features were primarily associated with earlier time lags relative to the linguistic features (**Fig. 2C**). A strong correlation was observed between the ISC values for component 1 and 2 and the prediction accuracies for the univariate models, particularly for WO and WS (**Fig. 3A**). Moreover, the ISC for component 1 and component 2 were significantly reduced when subtracting the predictions of each univariate model (**Fig. 3B, Table S2, Table S3**), with the acoustic features contributing more to ISC than the linguistic features, and WS more than WO (**Fig. 3B, Table S2, Table S3**).

**Fig 3.**
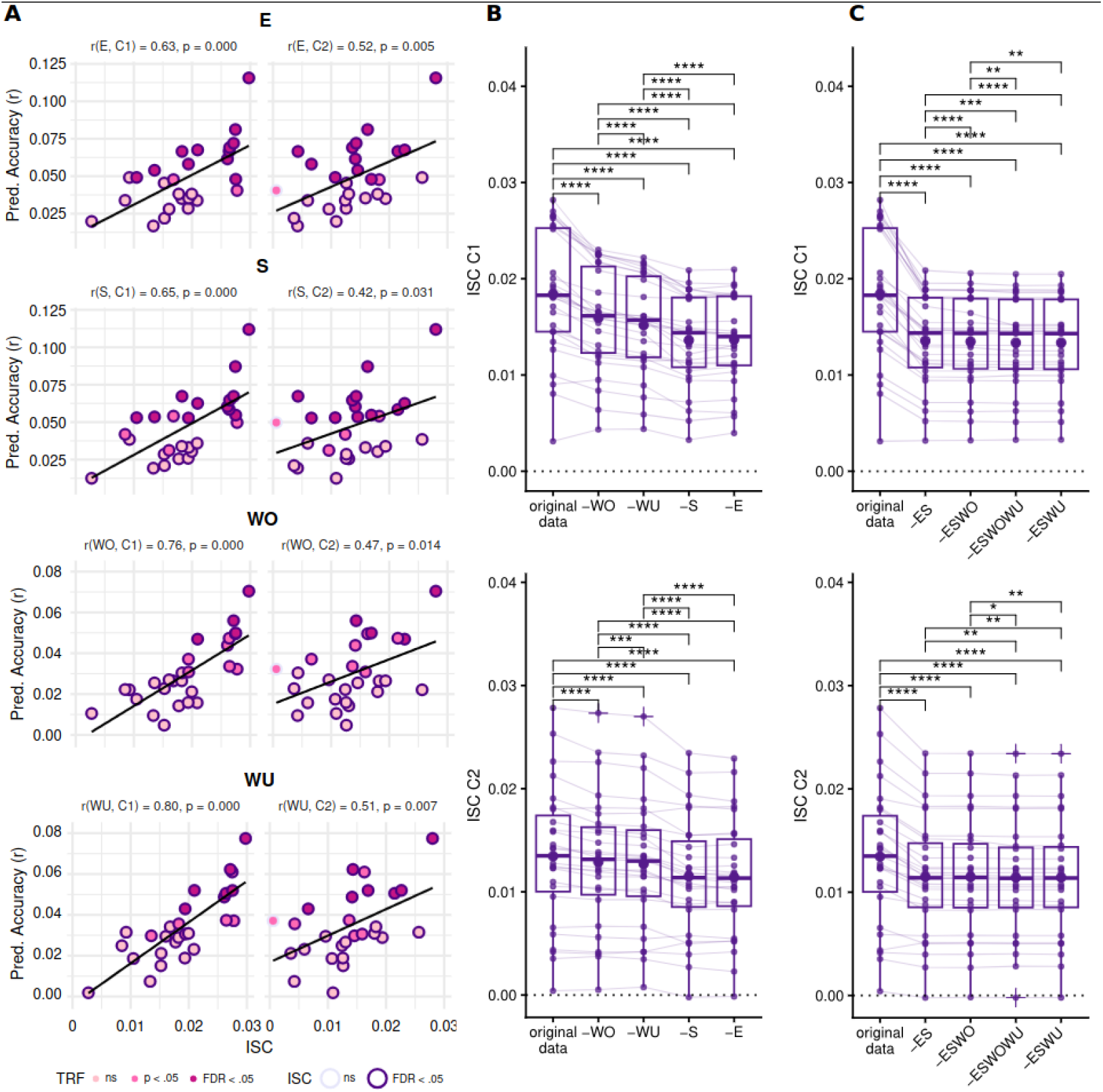
Speech features contribute to intersubject correlation. **A**. Correlation between the components that show the highest intersubject correlation (components 1 and 2, see **Fig S2** for component 3) and the average prediction accuracy for the univariate TRFs. Each point corresponds to a participant. Color edges indicate statistical significance for ISC: purple edge denotes significant EEG ISC (p < 0.05, FDR, corrected), grey edges non-significant values. Color fill indicates TRF statistical significance: dark pink marks a significant correlation between real and predicted EEG signal (p < 0.05, FDR corrected), light magenta denotes p < 0.05 without FDR correction, and light pink denotes p > 0.05. **B**. ISC computed over EEG residuals after subtracting the predicted activity of each univariate model. −E: without envelope TRF prediction, −S: without spectrogram TRF prediction, −WO: without word onset TRF prediction, −WS: without word surprisal prediction. Top: ISC for component 1. Bottom ISC for component 2. **C**. ISC computed over EEG residuals after subtracting the predicted activity of each multivariate model. −ES: without envelope and spectrogram TRF prediction, −ESWO:- without envelope, spectrogram and word onset TRF prediction, −ESWO:- without envelope, spectrogram and word onset TRF prediction, and −ESWOWS:- without envelope, spectrogram, word onset and word surprisal TRF prediction. Top: ISC for component 1. Bottom ISC for component 2. In each boxplot, the horizontal line within the box denotes the median and the dot represents the mean. Two-tailed Wilcoxon signed-rank test, (*) p < 0.05, (**) p < 0.01, (***) p < 0.001, and (****) p < 0.0001, FDR corrected.

### Intersubject correlation and speech tracking are positively modulated by attention

The purpose of Experiment 2 was to assess how attention influences ISC and whether attentive listening of narratives uniquely affects the linguistic contributions. We compared the ISC during attentive listening and distracted listening. As expected, ISC for the first three CorrCA components was reduced during the distracted condition (**Fig. 4A**, **Table S6**).

**Fig 4.**
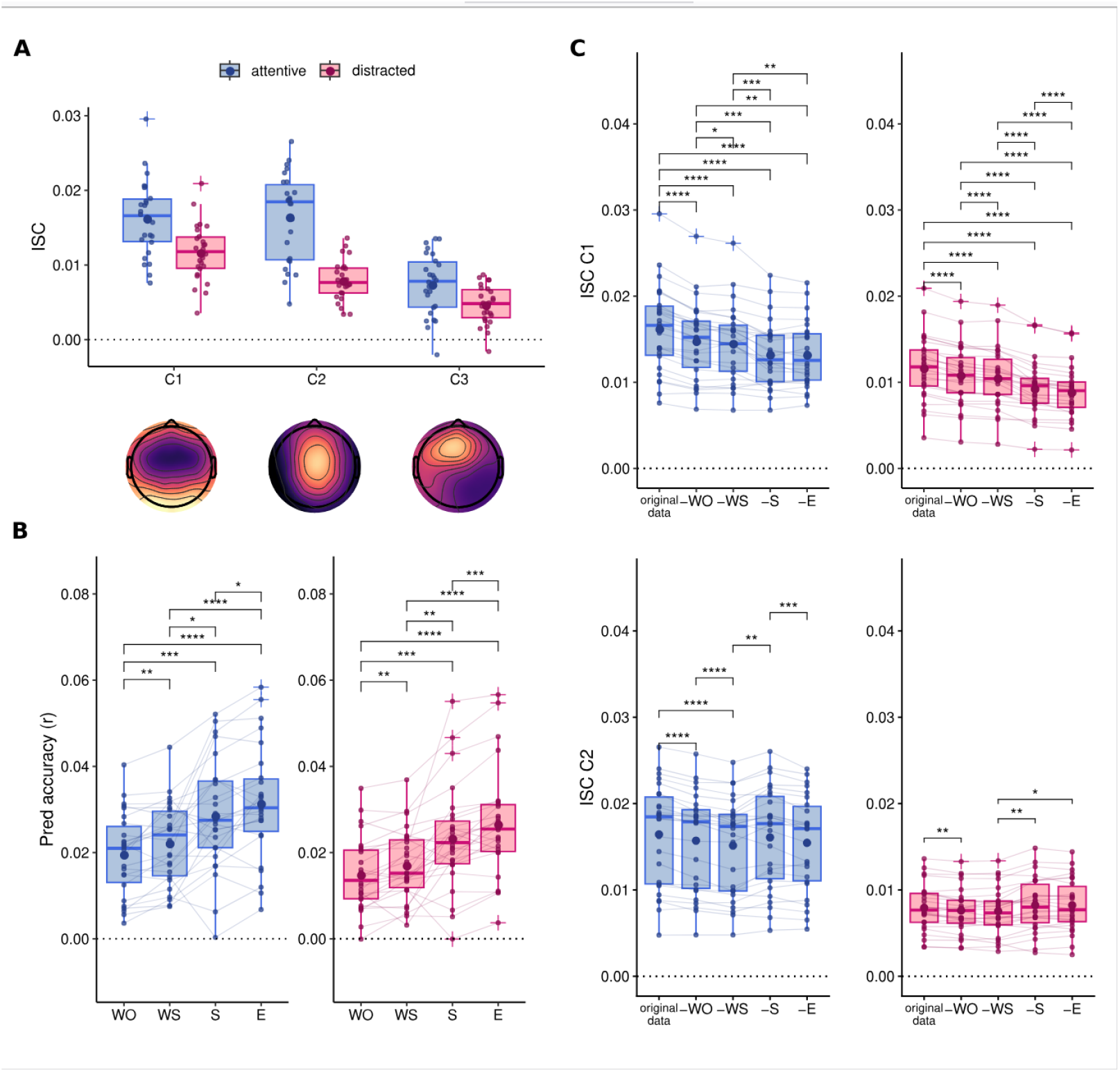
Speech features contribute to intersubject correlation while attentive and distracted listening of the narratives. **A**. ISC for the first three strongest components by attentional condition. Bottom. Forward model for the components. **B**. Prediction accuracy for each univariate model by condition. **C**. ISC computed over EEG residuals after subtracting the predicted activity of each univariate model for component 1 (top) and component 2 (bottom). In each boxplot, the horizontal line within the box denotes the median and the dot represents the mean. −E: without envelope TRF prediction, −S: without spectrogram TRF prediction, −WO: without word onset TRF prediction, −WS: without word surprisal prediction. Two-tailed Wilcoxon signed-rank tests, (*) p < 0.05, (**) p < 0.01, (***) p < 0.001 and (****) p < 0.0001, FDR corrected.

### Linguistic features contribute distinctively to the second correlated component only when attention is engaged

The prediction accuracy for each univariate model was higher during attentive listening as compared to distracted listening (**Fig. 4B, Fig. S5, Table S7**). Independently of the attentional condition, models for acoustic features explained a larger fraction of the EEG (**Fig 4B, Table S8**). Interestingly, the prediction accuracies for the word surprisal encoding models were better than for word onset when participants both attended and ignored the stories (**Fig 4B, Table S8**).

The TRF weights for the WO and WS obtained with the common ridge regression parameter across participants were assessed with a one-way paired sample t-test corrected for multiple comparisons using a cluster-level statistical permutation test. Both WO and WS TRFs showed more negative weights in left temporal and parietal electrodes during the attentive compared to the distracted condition in a time window ∼500 ms post word onset (WS_attended_ - WS_distracted_: t_sum_ = −177, p = 0.013, t = 456:638 ms, max effect size: d = −0.92, T7, t = 581 ms; WO_attended_ - WO_distracted_: t_sum_ = −159, p = 0.015, t = 472:613 ms, max effect size: d = −0.91, FC5, t = 503 ms) (**Fig. 5B**). In addition, WS TRFs were significantly more negative than WSr TRFs during attended (t_sum_ = −1133, p = 0.0005, t = 347:675 ms) and distracted conditions (t_sum_ = −921, p = 0.0005, t = 300:675 ms) for central parietal electrodes (**Fig. 5B**).

**Fig 5.**
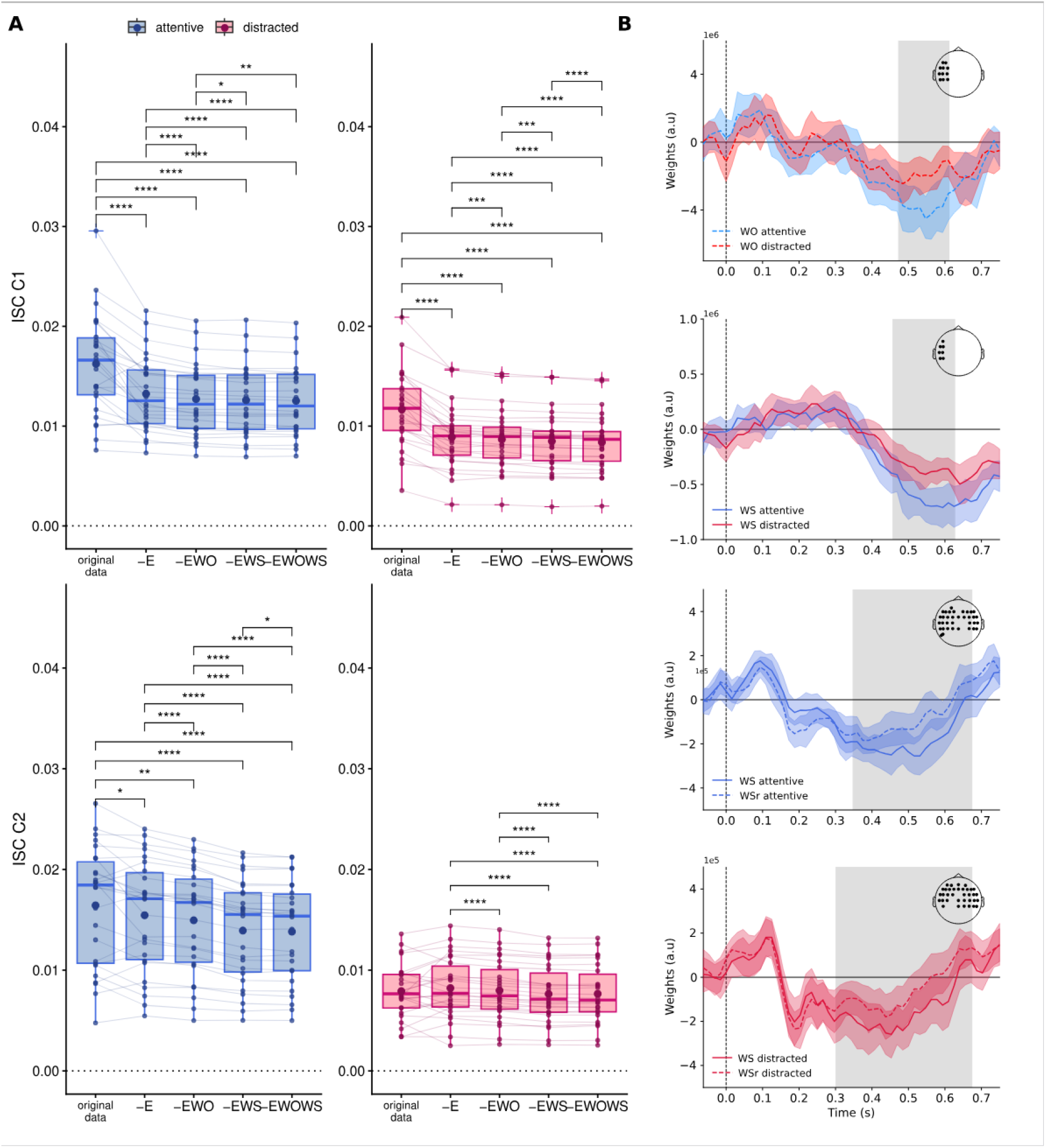
ISC during attentive speech listening is elicited by linguistic integration. **A.** ISC for original data during attentive and distracted speech listening, and computed over EEG residuals after subtracting the predicted activity of the multivariate model that showed the best predictions in both conditions (-EWO: without envelope and word onset model prediction, −EWS: without envelope and word surprisal model prediction). Top: ISC for component 1. Bottom ISC for component 2. In each boxplot, the horizontal line within the box denotes the median and the dot represents the mean. Two-tailed Wilcoxon signed-rank test, (*) p < 0.05, (**) p < 0.01, (***) p < 0.001 and (****) p < 0.0001, FDR corrected. **B.** TRF weights obtained with optimal λ across subjects and conditions. Grey shadings denote time span and inset shows electrodes that take part in significant clusters (WS: word surprisal, WSr: word surprisal values randomized within trials).

The predicted EEG by each univariate encoding model was individually subtracted from the original brain activity and the ISC was recomputed. The residual correlated components when projected to the 64-channel space had the same topographic distribution as the original data (**Fig. S4**). For the first component, the predicted activity of all univariate models contributed to ISC regardless of attentional condition, showing the same pattern observed during passive listening (**Fig. 4C top, Table S9**). In contrast, the ISC of the second component was largely unaffected by removal of the evoked activity predicted by the acoustic models, as neither comparison survived correction for multiple comparisons. Complementary Bayesian analyses provided only weak evidence for a difference between the original ISC and the ISC after removal of activity predicted by the envelope encoding model (BF₁₀ = 2.12), while providing modest evidence in favor of the null for the corresponding comparison for the spectrogram. Importantly, the contribution of the WO and WS models depended on attention: the activity predicted by the WS model influenced the ISC beyond WO only when participants were attending to the stimuli (**Fig. 4C bottom, Table S10**).

To account for correlations among speech features and to better assess the relationship between linguistic features, ISC, and attention, we constructed multivariate models comprising all combinations of the E, S, WO, and WS features. Pairwise comparisons between mTRFs showed a positive modulation of attention on their prediction accuracy (**Fig S6, Table S11**) and the models that showed the highest accuracies (EWO, EWS, and EWOWS) were selected to further explore the effects on the ISC (**Fig. S7**). Following the univariate models, removing the predicted activity by the EWS and EWO models caused a decrease in ISC for component 1 independently of the attentional condition with a higher contribution elicited by the models including WS (**Fig. 5A top, Table S12**). Crucially, models incorporating word surprisal provided the most accurate predictions of the ISC for the second component, but only when participants were actively attending to the stories (EWO vs. EWS: W = 350, df = 25, p < 0.0001). Moreover, the model combining the envelope with both linguistic features produced the highest decline in the ISC indexed by component 2 (**Fig. 5A bottom, Table S13**). Finally, the drop in ISC is specific of linguistic integration and is not produced by the mere addition of features as the same multivariate model with a randomized version of the WS feature (EWOWSr) explained less of the correlated activity than the EWS and EWOWS models (**Fig. S8**). Consistent with these findings, the contribution of word surprisal beyond speech acoustics to the ISC was also evident during passive listening. Although none of the mTRFs predicted the EEG response significantly better than the others **(Fig. S3)**, multivariate models including WS resulted in the largest reduction in ISC for components 1 and 2 **(Fig. 3C; Table S4; Table S5)**.

A key result of this analysis is that regressing out all explored features leads to only a modest reduction in intersubject correlation. Specifically, ISC for the first component decreased from 1.62 × 10⁻² to 1.25 × 10⁻², while ISC for the second component decreased from 1.64 × 10⁻² to 1.38 × 10⁻² (∼20%). This limited reduction suggests that a large fraction of activity elicited by the narrative and shared across subjects is driven by more complex responses or by features not examined in the current study.

## Discussion

The present study aimed to explore whether the EEG-correlated responses produced when participants listen to the same narratives are elicited by the processing of basic acoustic information of the speech signal or higher linguistic representations related to the content of speech, and how attention to the narratives influences these responses. For this, we built subject-level encoding models from the envelope, the spectrogram, the word onset, and word surprisal, and assessed their contributions to the ISC elicited during passive, attended, and distracted speech processing.

### ISC captures complex features of the narrative stimulus

The most salient observation from our analysis is that only a fraction of the ISC can be explained by responses to features traditionally used in the analysis of EEG during continuous speech (Broderick et al., 2018, 2019, 2022; Dou et al., 2024; Gillis et al., 2021, 2023; Shyanthony R. Synigal et al., 2023; Teoh et al., 2022). A key limitation of the TRF approach is its dependence on a predefined set of stimulus features, as well as its restriction to linear mappings within a limited temporal window (here, 750 ms). As a consequence, more complex or delayed responses to the narratives are not captured by the TRF formalism. In contrast, ISC captures any stimulus-related responses, provided they are shared across subjects. The unexplained variance by the TRFs may reflect neural integration across multiple timescales spanning sub-lexical and lexical features, higher-order linguistic units such as phrases and constituents (Gwilliams et al. 2024), and the resulting conceptual abstractions evoked by the stories. An alternative, however, may be to think of responses in the broader view of arousal fluctuations elicited by the narratives. Recent work has shown that significant variance in the EEG during auditory narratives is predicted by physiological signals (e.g. heart rate, breathing, pupil dilation) as well as incidental movements (head and eye movement), which themself synchronize across listeners during narrative engagement (Madsen et al., n.d.; Madsen & Parra, 2024). In this view, ISC may capture indirect, arousal-mediated effects of narrative processing.

### Encoding models and attention effects

We observed enhanced speech cortical tracking for both acoustic and linguistic features when participants attended to the stories as reflected by higher prediction accuracies in both univariate and multivariate models. This is consistent with research assessing the effects of attention on neural tracking of the envelope (Rosenkranz et al., 2021; Vanthornhout et al., 2019), the spectrogram (Teoh et al., 2022), semantic information (Broderick et al., 2018) and lexical surprise (Shyanthony R. Synigal et al., 2023). The weights of the envelope encoding models exhibited topographic and temporal patterns closely resembling classical auditory N1/P2 event-related potentials (ERP) (Čeponienė et al., 2008) as well as temporal response functions reported in previous studies (Di Liberto et al., 2015; Mesik & Wojtczak, 2023) (**Fig. 2C**). Similarly, the word surprisal model weights were consistent with previous research (Dou et al., 2024) and reminiscent of the N400 ERP (Kutas & Hillyard, 1980), for which amplitude is modulated by, among other properties, word context (Kutas & Federmeier, 2014) (**Fig. 2C**, **Fig. 5C**). In predictive models of language processing, surprisal is considered to reflect the amount of information gained when prior expectations about upcoming words are updated, indexing the cognitive effort required for comprehension (Hale, 2001; Levy, 2008). Accordingly, word surprisal has been shown to be a good predictor of the N400 amplitude (Frank et al., 2015; Michaelov et al., 2024; Michaelov & Bergen, 2020; Szewczyk & Federmeier, 2022). In the present study, contrasting the word surprisal encoding models with a shuffled version of this representation yielded more negative weights over central-parietal electrodes within a time window consistent with that reported in the N400 literature, further supporting the notion that the model captures context-dependent linguistic information (**Fig. 5C**). Interestingly, this effect was also observed for unattended stimuli suggesting that some degree of lexical processing was occurring outside of conscious awareness. Consistent with this view, semantic access without awareness has been reported in auditory word paradigms during sleep (Andrillon et al., 2016) and coma (Rämä et al., 2010). However, neural evidence for unconscious semantic integration beyond isolated words is scarce (Mudrik & Deouell, 2022). In this context, because the attentional manipulation was not accompanied by an objective behavioral measure, it is difficult to assess participants’ compliance with the task and, consequently, the extent to which attention was successfully diverted from the auditory stimuli. Therefore, the present finding may alternatively reflect imperfect suppression or filtering of the speech signal during the distracted condition rather than genuine unconscious integration of context-dependent meaning.

### Acoustic information explains a larger fraction of the ISC elicited by narratives

ISC was elicited during passive listening as well as during attended and distracted speech processing, with values exceeding chance for the first and second correlated components. Moreover, the resulting scalp topographies were consistent with those reported in previous studies (Cohen and Parra 2016; Dmochowski et al. 2012; Ki, Kelly, and Parra 2016; Petroni et al. 2018; Rosenkranz et al. 2021). During passive listening and during both attentional conditions in Experiment 2, acoustic information captured by the spectrogram and the envelope contributed more strongly to ISC than word onset or word surprisal features. This pattern can be partially attributed to differences in feature representation: acoustic features yield dense, continuous representations, whereas the linguistic features considered here are inherently sparse and therefore tend to produce lower prediction accuracies (Chalehchaleh et al., 2025). Nevertheless, despite this methodological disadvantage, linguistic features still made a meaningful contribution to ISC, suggesting that their true impact may be underestimated by the present approach.

### Linguistic integration produces specific shared evoked activity only when attending to the stories

In previous work, the ISC of the EEG has been postulated as a marker of engagement during film viewing (Cohen & Parra, 2016; Dmochowski et al., 2012; Poulsen et al., 2017) and music listening (Madsen et al. 2019). In agreement, in our study ISC was positively modulated by attention, with a particularly stronger effect on the correlated activity indexed by the second correlated component. In addition, a distinctive pattern of linguistic and acoustic contributions to ISC emerged as a function of the attentional conditions. The first ISC component was driven by acoustic features independently of attention, as a greater decrease in ISC was observed when removing the EEG predictions by models built from acoustic features compared to linguistic features during attended, distracted and passive listening. In contrast, the ISC indexed by the second correlated component was driven by linguistic contributions only during the attentive condition and during passive listening, as supported for both univariate and multivariate prediction subtractions. Ki and collaborators (Ki et al. 2016) have previously shown an attentional modulation of the ISC reflected by this component for audiovisual narratives, with a smaller effect for scrambled narratives and no effect of attention when the narrative was presented in a foreign language. Here we provide evidence of a specific contribution of linguistic information on the second component ISC beyond word onset, as the multivariate models based on word surprisal explained more of the intersubject correlation than equivalent models based on word segmentation. The word surprisal representations reflect the probability of that word based on multiple types of linguistic information. BERT models learn structural properties of language such as syntax, but also semantic roles, and some types of world knowledge (Rogers et al., 2020), thus we summarize this into the term ‘linguistic integration’. Therefore, our results support the hypothesis that the activity indexed by the second correlated component is probably capturing shared brain dynamics related to linguistic integration while the first correlated component is elicited mainly by the low-level sensory processing of speech, being both sensitive to attention. In fact, these two processes are closely related, as a directional effect in which lexical information influences the encoding of acoustic features has been described (Broderick et al., 2019; Heilbron et al., 2022). In addition, our results show that even under passive listening, a scenario without task-related demands which closely reflects natural speech processing, robust encoding models can be built and linguistic information contributes to the correlated neural response captured by the second component beyond what can be explained by acoustic or segmentation features alone.

### Considerations for studies of unresponsive patients

Unlike experiments where isolated words or short sentences are presented to participants, experiments with narrative stimuli allow probing of the brain in a more naturalistic and engaging way, providing better information on the mechanisms behind the perception of language (Hamilton & Huth, 2020; Sonkusare et al., 2019). Moreover, participants can be tested with passive paradigms, which allows the comparison of language processing between healthy controls and pathological populations for which following instructions or providing verbal or motor outputs is not possible (Sokoliuk, Degano, Melloni, et al., 2021). In the case of patients with disorders of consciousness (DoC), where prognostic information is fundamental to guide the decisions of caregivers regarding treatment (Russell et al., 2024), residual language capacities are associated with better outcomes (Coleman et al., 2009; Gui et al., 2020; Sokoliuk, Degano, Banellis, et al., 2021). Therefore, developing tools that are easy to implement and that assess with granularity the level of speech processing in patients with DoC is of extreme value. Importantly, if the comprehension of complex meaning requires awareness (Mudrik & Deouell, 2022; Rabagliati et al., 2018), having markers of linguistic integration would provide support for the conscious processing of information in a patient, pointing to states of cognitive motor dissociation (Bodien et al., 2024; Claassen et al., 2024) or covert cortical processing (Edlow et al., 2017), which are considered prognostically promising.

Whether ISC is driven by basic acoustic stimulus features or language processing is important to interpret some recent clinical results. Two studies have assessed EEG responses of patients with disorders of consciousness (DoC) when exposed to auditory narratives, showing that some patients present brain activity patterns correlated to the evoked responses of healthy controls (Laforge et al., 2020). In addition, ISC is generally decreased in these patients, and the strength of the ISC differs during forward and backward speech (Iotzov et al. 2017), a phenomenon that may relate to the clinical diagnosis. Were those patients encoding auditory features, segmenting speech and/or integrating complex meaning?. Identifying which speech properties elicit these common brain responses is crucial to understanding what language capabilities, including conscious comprehension, are retained by a DoC patient who exhibits ISC. In light of our results, we propose that combining ISC with encoding models provides a principled framework to address this question and to characterize the language capabilities retained by patients who exhibit ISC. Importantly, our results show that a passive listening scenario is sufficient to detect acoustic and linguistic speech tracking at the subject level, rendering this approach highly appropriate as a bedside examination to determine the depth of speech processing in DoC patients. In this line, we believe our work opens a clear avenue for a feasible and informative assessment, and we offer some recommendations for its design.

Firstly, selecting a well-thought-out stimulus is of paramount importance. Stories should be engaging, use broadly familiar vocabulary, at least 15 minutes long (Crosse et al., 2021), and preferably conveyed by a unique speaker to minimize acoustic variability. Some research has used forward and backward language to compare language processing in DoC patients (Fernández-Espejo et al., 2008; Iotzov et al., 2017), we argue that an optimal control would be to use a story in an unfamiliar language. Although reverse speech can convey information on language-independent auditory processing (Fernández-Espejo et al., 2008), it is questionable whether it has the same statistical properties as actual speech. Brain response in a cohort of healthy controls exposed to the story should be recorded in two attentional conditions counterbalanced across participants. Comprehension questions should be posed at the end of the attentive condition, and the distracted condition should be demanding enough to prevent participants from also engaging with the story. ISC and encoding models for acoustic and linguistic features would be computed, and it should be verified that at least some of the ISC is conveyed by these representations. A patient would be presented with the same stimuli and be asked to actively listen to the story. Encoding models for the different features would provide specific information on whether the patient is processing sensory information and also linguistic information. If more sophisticated linguistic models are not available, combining acoustic features with word onset predictors can still provide informative markers of speech segmentation. The discreet phenomenology of speech does not occur with an unfamiliar language (Ding et al., 2016), as it requires knowledge about the transition probabilities between speech sounds and their correspondence to word boundaries, a mapping acquired through statistical learning (Erickson & Thiessen, 2015). In addition, word tracking is not elicited during REM and non-REM sleep (Makov et al., 2017). Therefore, comparing the encoding models for word onset during native and unfamiliar language would be of interest. Finally, the patient’s brain response to the narratives should be correlated to the healthy cohort response during attended and distracted speech listening. If the patient shows a greater correlation with the attended than the distracted responses, while also showing that the evoked activity predicted by linguistic models affects the ISC indexed by the second correlated component, then we could infer language processing beyond simple acoustics. Moreover, given that most of the ISC is not explained by the features assessed in this work, a patient that correlates highly to the healthy cohort during the attentive condition is probably also responding as healthy participants to other sources of information, or integrating the stimuli in non-linear ways, suggesting highly conserved cognitive capabilities such as executive functions, attention and working memory.

In summary, the complementary information provided by the ISC and the model-based predictions would enable evidence-based inferences on the level of residual language processing in DoC patients.

### Limitations

Some aspects of our study should be considered when interpreting the results as well as to improve future work. First, the attentional conditions in Experiment 2 were not counterbalanced across participants and could have influenced our results. Specifically, because the distracted condition always involved a second exposure to the stories, we cannot entirely rule out contributions from familiarity or fatigue effects to the observed reductions in ISC and speech-feature encoding. Nevertheless, our findings are consistent with previous evidence showing that ISC and cortical tracking of speech are reduced when attention is diverted. Speech features continued to be encoded during the distracted condition, and this neural response was shared across individuals. However, we cannot rule out the possibility that this response resulted from imperfect filtering of the auditory stimuli. A replication implementing an objective behavioral measure of task following would further strengthen our interpretations by providing a direct assessment of whether attention was successfully diverted from the auditory stimuli. Finally, in Experiment 2 participants listened to several stories, some of which featured multiple speakers. This variability in the attended-distracted experiment meant that our models had to generalize to more dissimilar stimuli compared to Experiment 1 which may help explain why the multivariate models that showed greater prediction accuracies differed between both experiments.

Experiments 1 and 2 differed substantially in several respects, including stimulus material and speaker identity, recording setups, preprocessing pipelines, and experimental structure. In particular, Experiment 1 involved a single story and speaker in a passive-listening paradigm, whereas Experiment 2 used multiple stories and speakers and included attentive and distracted conditions. These differences preclude direct comparisons of the magnitude of ISC or model performance across experiments. Nevertheless, a similar qualitative pattern emerged across the datasets: the correlated activity of the second component during passive listening, as well as during attentive listening, was associated not only with acoustic features but also with higher-level linguistic features, specifically lexical surprise. We therefore interpret this convergence across these distinct datasets as evidence for the robustness and generalizability of our approach.

### Conclusions and future directions

We show that a proportion of the intersubject correlation evoked by listening to narrative stimuli is produced by both acoustic and linguistic content present in the speech signal. The increases in ISC produced by engaging with the stimuli are driven by an enhancement in the neural representations of both types of information, with a specific effect of linguistic integration on the second correlated component when attending to the stories. Intriguingly, the explored features explain a small amount of the overall ISC and future work should address which other speech properties are at play. Importantly, we show that narrative stimuli can be used to capture acoustic and linguistic speech tracking at the subject level, even in a passive listening scenario, rendering this approach highly appropriate as a bedside examination to determine low-level and high-level feature encoding as well as broader neural responses reflecting general engagement with the stimuli.

Overall, our results suggest that a proportion of the ISC of the EEG arises from the integration of multiple levels of information present in speech and propose that ISC, together with encoding models, could provide meaningful information on language processing in unresponsive patients.

## Ethics approval and consent to participate

Experiment 1 was approved by the STEM ethics committee of the University of Birmingham, England. Experiment 2 was approved by the Institutional Review Boards of the City University of New York. All subjects provided written informed consent before the experiment.

## Availability of data and materials

The datasets generated and analyzed during the current study are available from the corresponding author upon reasonable request. The code used for preprocessing and analysis is openly available at https://github.com/Em-Fl/TRF_ISC to facilitate the reproduction of the analyses.

## Authors Contributions

Conceptualization: EF and JS. Methodology: EF and AC. Formal analysis: EF. Visualization: EF. Resources: JS, DC, JM, LCP. Writing - original draft: EF. Writing - review & editing: EF, AC, JVL, DC, JM, LCP, JS. Supervision: JS. Funding acquisition: EF, AC, JVL, JS

## Supplementary materials

**Fig S1.**
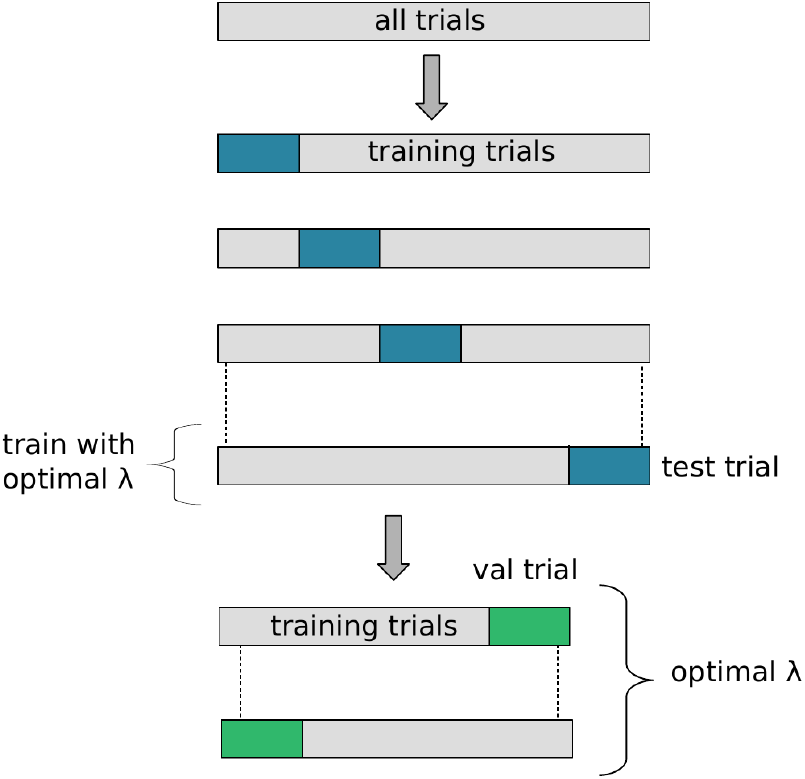
Leave-one-out nested cross-validation to select the ridge parameters. For each subject, we iteratively selected one trial to test the TRF (in blue) and used the remaining trials to perform a leave-one-out cross-validation to obtain the ridge parameter across channels and folds that maximized the correlation between the actual response and the predicted response. The n-1 trials used in cross-validation were used to train the model with the optimal lambda, and the model was then tested on the left-out trial. This iterative process yielded one Pearson correlation coefficient per trial and subject, given by the average correlation across all channels.

**Table 1:** Passive listening: prediction accuracy for univariate models.

| x | y | W | p | padj<sup>1</sup> |
| --- | --- | --- | --- | --- |
| WO | WS | 31 | 3.52e-05 | **7.04e-05** |
| WO | S | 1 | 2.98e-08 | **1.79e-07** |
| WO | E | 4 | 1.04e-07 | **5.20e-07** |
| WS | S | 6 | 2.09e-07 | **6.27e-07** |
| WS | E | 6 | 2.09e-07 | **6.27e-07** |
| S | E | 138 | 2.29e-01 | 2.29e-01 |

<sup>1</sup> FDR corrected. Bold indicates $p < 0.05$
Wilcoxon two-sided signed-rank test results for encoding models' prediction accuracy. WO: word onset, WS: word surprisal, S: spectrogram, E: envelope.

**Fig S2.**
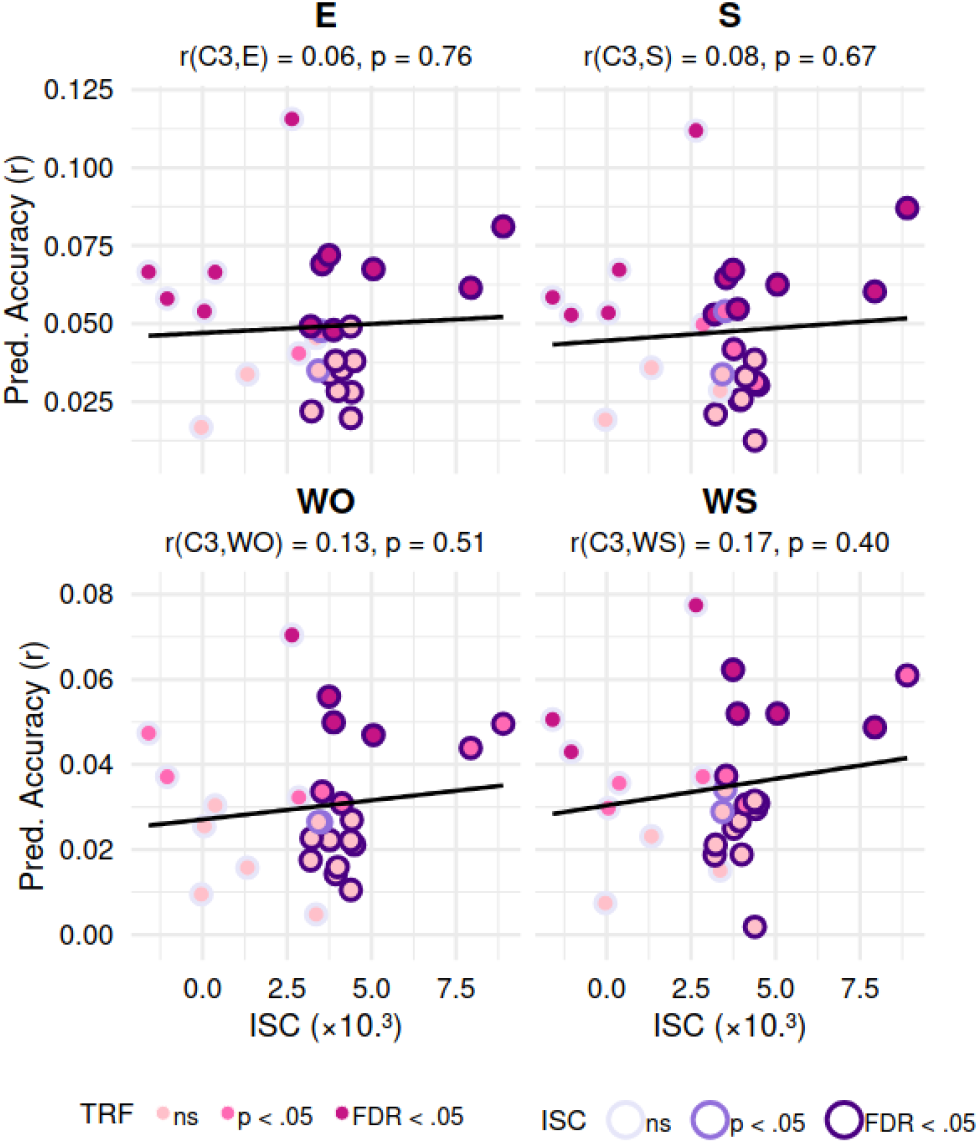
Correlation between the ISC for component 3 and the average prediction accuracy for the univariate TRFs. Each point corresponds to a participant. Color edges indicate statistical significance for ISC: purple edge denotes significant EEG ISC(p < 0.05, FDR, corrected), lila edges indicate p<0.05 without FDR correction, and grey edges indicate non-significant values. Color fill indicates TRF statistical significance: dark pink marks a significant correlation between real and predicted EEG signal (p < 0.05, FDR corrected), light magenta denotes p<0.05 without FDR correction, and light pink denotes p>0.05. WO: word onset, WS: word surprisal, S: spectrogram, E: envelope.

**Table S2:** Passive listening: univariate contributions to ISC component 1.

| x | y | W | p | p<sub>adj</sub><sup>1</sup> |
| --- | --- | --- | --- | --- |
| original data | -WO | 374 | 1.04e-07 | **3.12e-07** |
| original data | -WS | 376 | 4.47e-08 | **1.79e-07** |
| original data | -S | 377 | 2.98e-08 | **1.49e-07** |
| original data | -E | 377 | 2.98e-08 | **1.49e-07** |
| -WO | -WS | 377 | 2.98e-08 | **1.49e-07** |
| -WO | -S | 378 | 1.49e-08 | **1.49e-07** |
| -WO | -E | 377 | 2.98e-08 | **1.49e-07** |
| -WS | -S | 377 | 2.98e-08 | **1.49e-07** |
| -WS | -E | 371 | 2.83e-07 | **5.66e-07** |
| -S | -E | 156 | 4.41e-01 | 4.41e-01 |

<sup>1</sup>FDR corrected. Bold indicates $p < 0.05$
Wilcoxon two-sided signed-rank test results for ISC of component one after encoding model subtraction. WO: word onset, WS: word surprisal, S: spectrogram, E: envelope. The (-) symbol indicates that the predicted activity by the encoding model was subtracted from the EEG response.

**Table S3:** Passive listening: univariate contributions to ISC component 2.

| x | y | W | p | p<sub>adj</sub><sup>1</sup> |
| --- | --- | --- | --- | --- |
| original data | -WO | 359 | 4.57e-06 | **1.37e-05** |
| original data | -WS | 362 | 2.52e-06 | **1.37e-05** |
| original data | -S | 374 | 1.04e-07 | **1.04e-06** |
| original data | -E | 371 | 2.83e-07 | **2.55e-06** |
| -WO | -WS | 328 | 4.27e-04 | **8.54e-04** |
| -WO | -S | 362 | 2.52e-06 | **1.37e-05** |
| -WO | -E | 362 | 2.52e-06 | **1.37e-05** |
| -WS | -S | 360 | 3.77e-06 | **1.37e-05** |
| -WS | -E | 360 | 3.77e-06 | **1.37e-05** |
| -S | -E | 239 | 2.39e-01 | 2.39e-01 |

<sup>1</sup> FDR corrected. Bold indicates $p < 0.05$
Wilcoxon two-sided signed-rank test results for ISC of component two after encoding model subtraction. WO: word onset, WS: word surprisal, S: spectrogram, E: envelope. The (-) symbol indicates that the predicted activity by the encoding model was subtracted from the EEG response.

**Table S4:** Passive listening: multivariate contributions to ISC component 1.

| x | y | W | p | p<sub>adj</sub><sup>1</sup> |
| --- | --- | --- | --- | --- |
| original data | -ES | 377 | 2.98e-08 | **2.09e-07** |
| original data | -ESWO | 377 | 2.98e-08 | **2.09e-07** |
| original data | -ESWOWS | 377 | 2.98e-08 | **2.09e-07** |
| original data | -ESWS | 377 | 2.98e-08 | **2.09e-07** |
| -ES | -ESWO | 368 | 6.41e-07 | **3.85e-06** |
| -ES | -ESWOWS | 349 | 2.59e-05 | **1.04e-04** |
| -ES | -ESWS | 351 | 1.88e-05 | **9.40e-05** |
| -ESWO | -ESWOWS | 317 | 1.00e-03 | **3.00e-03** |
| -ESWO | -ESWS | 320 | 1.00e-03 | **3.00e-03** |
| -ESWOWS | -ESWS | 155 | 4.27e-01 | 4.27e-01 |

<sup>1</sup>FDR corrected. Bold indicates $p < 0.05$
Wilcoxon two-sided signed-rank test results for ISC of component one after encoding models subtraction. ES: envelope and spectrogram model, ESWO: envelope, spectrogram, and word onset model, ESWS: envelope, spectrogram, and word surprisal model, ESWOWS: envelope, spectrogram, word onset, and word surprisal model. The (-) symbol indicates that the predicted activity by the encoding model was subtracted from the EEG response before computing the ISC.

**Table S5:**
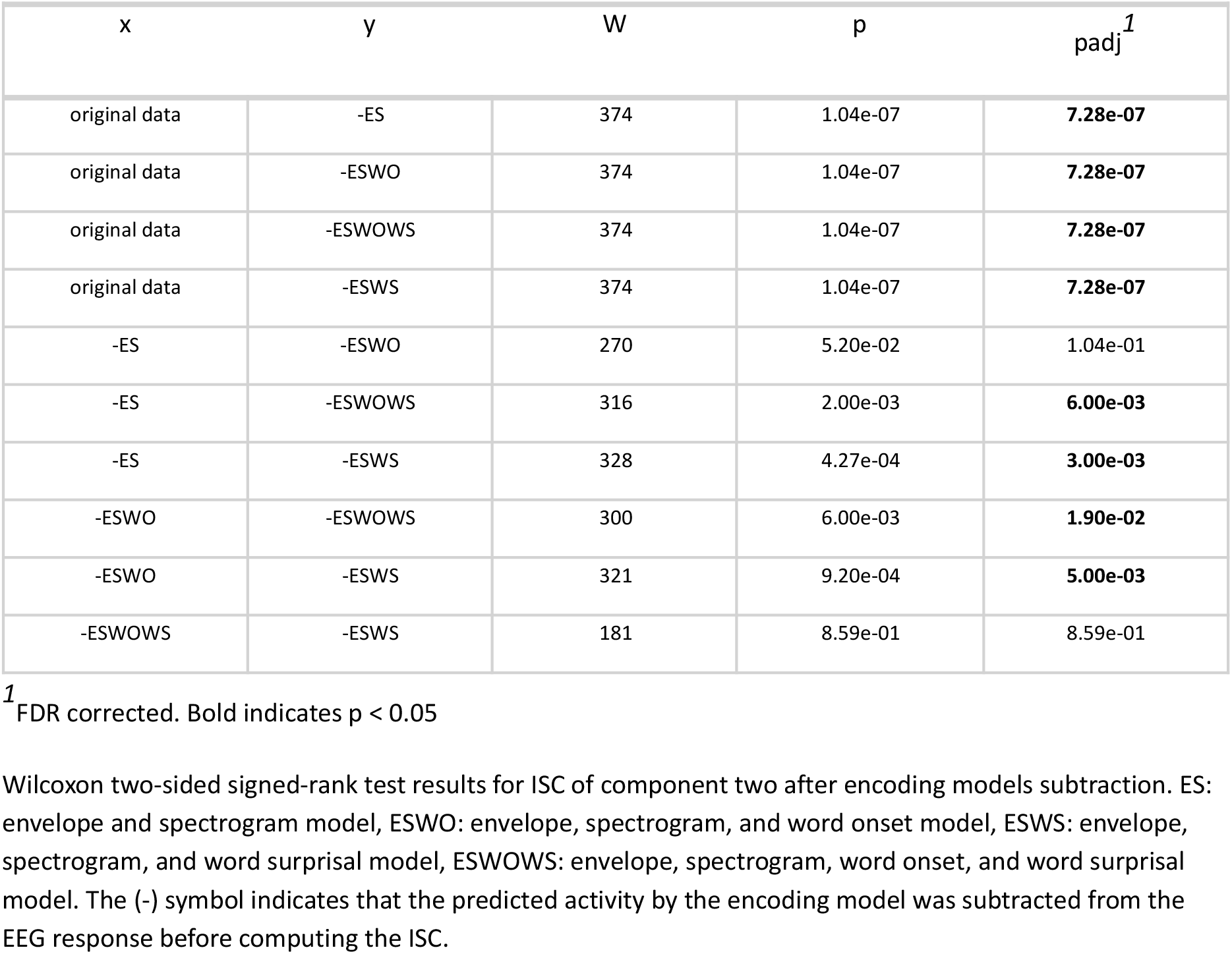
Passive listening: multivariate contributions to ISC component 2.

| x | y | W | p | p<sub>adj</sub><sup>1</sup> |
| --- | --- | --- | --- | --- |
| original data | -ES | 374 | 1.04e-07 | **7.28e-07** |
| original data | -ESWO | 374 | 1.04e-07 | **7.28e-07** |
| original data | -ESWOWS | 374 | 1.04e-07 | **7.28e-07** |
| original data | -ESWS | 374 | 1.04e-07 | **7.28e-07** |
| -ES | -ESWO | 270 | 5.20e-02 | 1.04e-01 |
| -ES | -ESWOWS | 316 | 2.00e-03 | **6.00e-03** |
| -ES | -ESWS | 328 | 4.27e-04 | **3.00e-03** |
| -ESWO | -ESWOWS | 300 | 6.00e-03 | **1.90e-02** |
| -ESWO | -ESWS | 321 | 9.20e-04 | **5.00e-03** |
| -ESWOWS | -ESWS | 181 | 8.59e-01 | 8.59e-01 |

<sup>1</sup> FDR corrected. Bold indicates $p < 0.05$
Wilcoxon two-sided signed-rank test results for ISC of component two after encoding models subtraction. ES: envelope and spectrogram model, ESWO: envelope, spectrogram, and word onset model, ESWS: envelope, spectrogram, and word surprisal model, ESWOWS: envelope, spectrogram, word onset, and word surprisal model. The (-) symbol indicates that the predicted activity by the encoding model was subtracted from the EEG response before computing the ISC.

**Fig S3.**
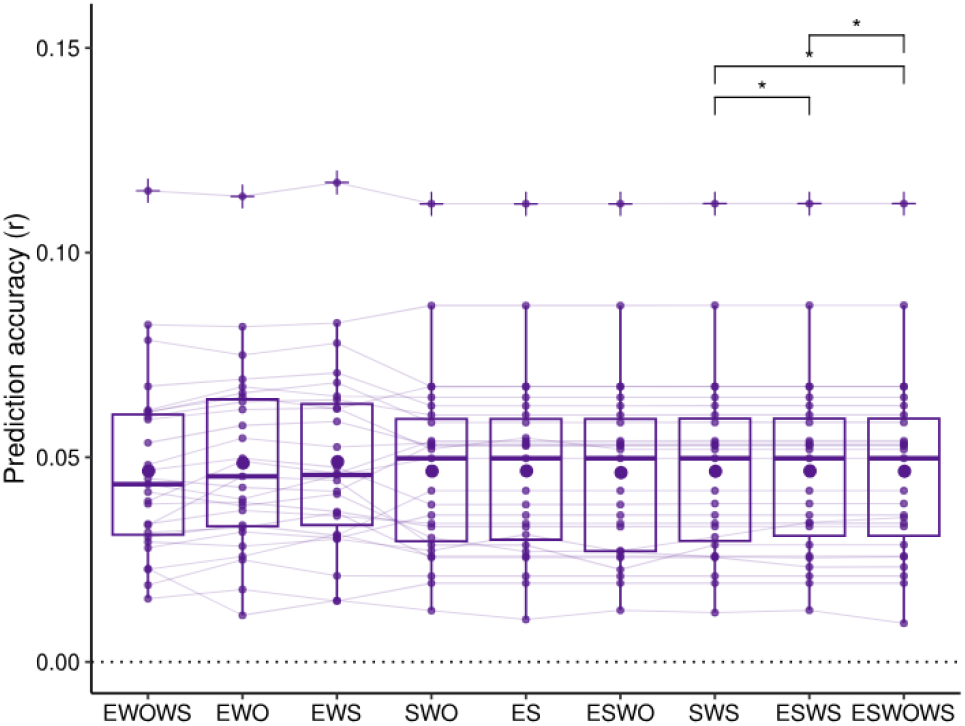
Passive listening: prediction accuracies for multivariate models. In each boxplot, the horizontal line within the box denotes the median and the dot represents the mean. E: envelope, S: spectrogram, WS: word surprisal, WO: word onset. Significant differences were only found between ESWS and SWS (W = 54, p = 0.023), SWS and ESWOWS (W = 52, p = 0.019), and between ESWS and ESWOWS (W = 48, p = 0.012). All other paired comparisons resulted in p-values ≥ 0.068.

**Table S6:** Attention effects on ISC.

| x | y | W | p | padj<sup>1</sup> |
| --- | --- | --- | --- | --- |
| attentive.C1 | distracted.C1 | 346 | 2.98e-07 | **5.96e-07** |
| attentive.C2 | distracted.C2 | 351 | 2.98e-08 | **8.94e-08** |
| attentive.C3 | distracted.C3 | 316 | 1.26e-04 | **1.26e-04** |

<sup>1</sup> FDR corrected. Bold indicates $p < 0.05$
Wilcoxon two-sided signed-rank test results for each ISC component during attentive and distracted conditions.

**Fig S4.**
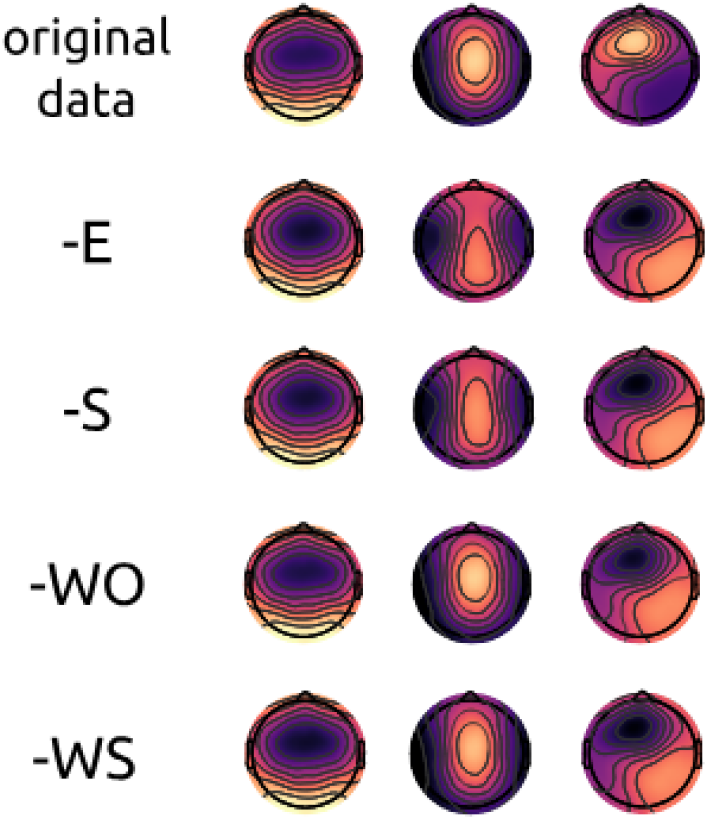
Forward model projections of the correlated components into sensor space for the original data, after subtraction of the envelope (-E), the spectrogram (-S), the word onset (-WO), and the word surprisal (-WS) encoding models’ predictions.

**Table S7:**
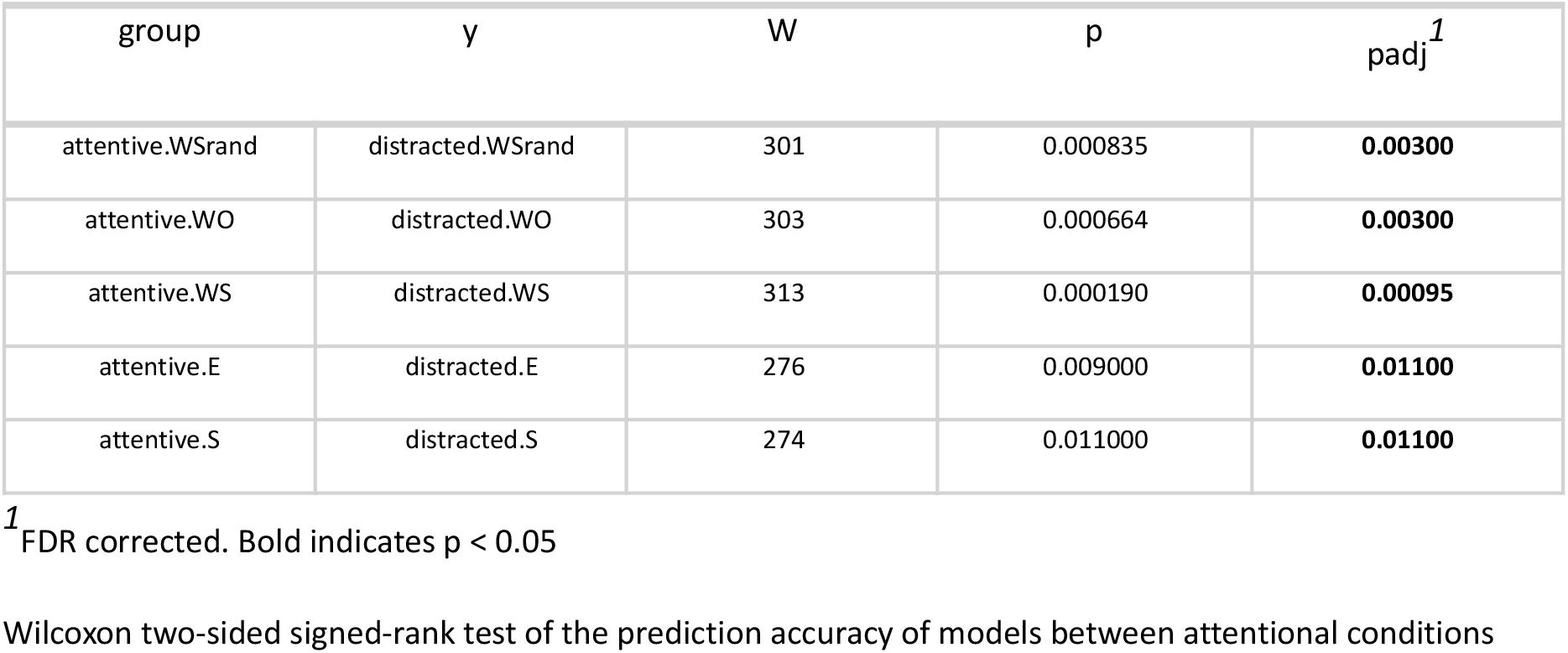
Attention effects on the prediction accuracies of the univariate TRFs.

| group | y | W | p | p<sub>adj</sub><sup>1</sup> |
| --- | --- | --- | --- | --- |
| attentive.WSrand | distracted.WSrand | 301 | 0.000835 | **0.00300** |
| attentive.WO | distracted.WO | 303 | 0.000664 | **0.00300** |
| attentive.WS | distracted.WS | 313 | 0.000190 | **0.00095** |
| attentive.E | distracted.E | 276 | 0.009000 | **0.01100** |
| attentive.S | distracted.S | 274 | 0.011000 | **0.01100** |

<sup>1</sup>FDR corrected. Bold indicates $p < 0.05$
Wilcoxon two-sided signed-rank test of the prediction accuracy of models between attentional conditions

**Fig S5.**
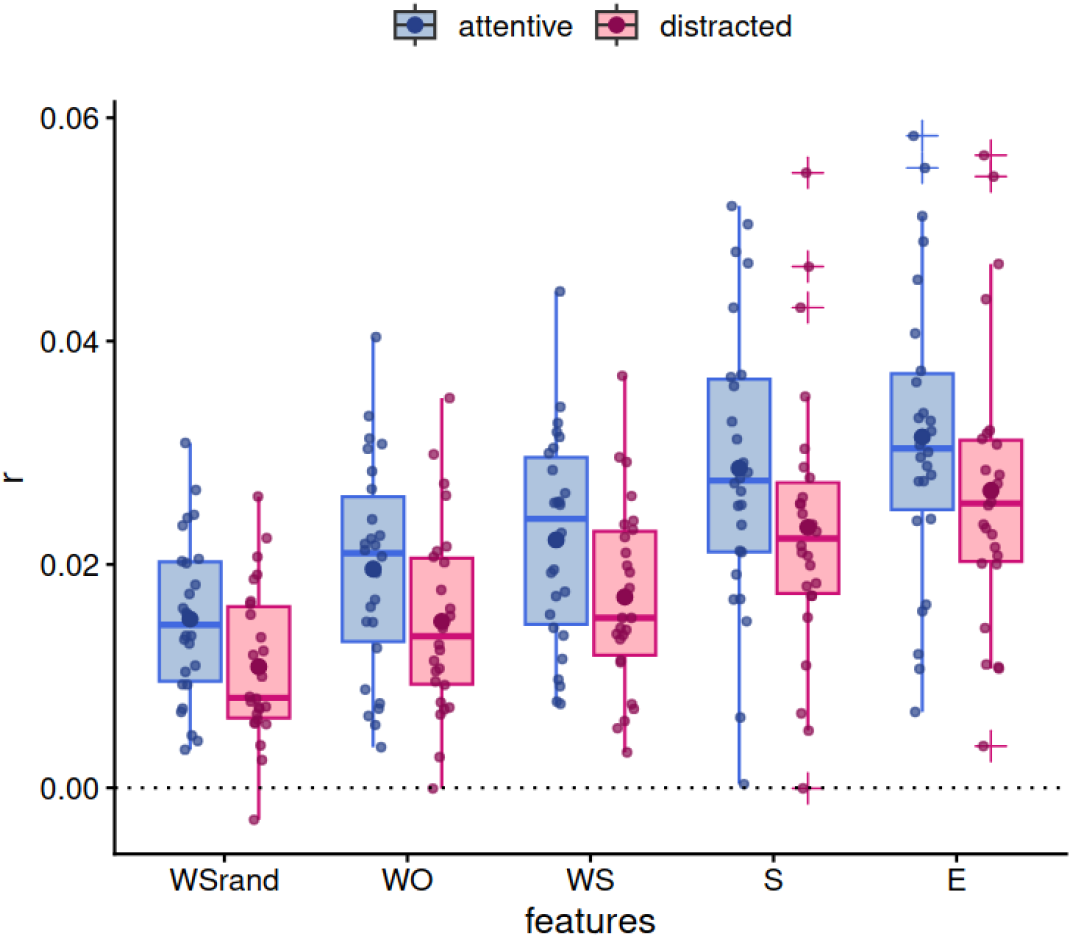
Prediction accuracies for univariate models during attentive and distracted conditions. In each boxplot, the horizontal line within the box denotes the median and the dot represents the mean. Envelope (E), spectrogram (S), word surprisal (WS), word onset (WO), and randomized word surprisal (WSrand). Attending to the stories results in higher prediction accuracies for all models.

**Table S8:** Prediction accuracies for univariate TRFs within attentional conditions.

| attentive |  |  |  |  | distracted |  |  |
| --- | --- | --- | --- | --- | --- | --- | --- |
| group 1 | group 2 | W | p | padj<sup>1</sup> | W | p | padj<sup>1</sup> |
| WO | WS | 57 | 2.00e-03 | **5.00e-03** | 60 | 2.00e-03 | **2.00e-03** |
| WO | S | 32 | 8.16e-05 | **3.26e-04** | 25 | 2.69e-05 | **1.08e-04** |
| WO | E | 4 | 2.09e-07 | **1.25e-06** | 2 | 8.94e-08 | **5.36e-07** |
| WS | S | 69 | 6.00e-03 | **1.00e-02** | 51 | 9.35e-04 | **2.00e-03** |
| WS | E | 18 | 7.54e-06 | **3.77e-05** | 13 | 2.62e-06 | **1.31e-05** |
| S | E | 76 | 1.00e-02 | **1.00e-02** | 33 | 9.45e-05 | **2.84e-04** |

<sup>1</sup>FDR corrected. Bold indicates $p < 0.05$
Wilcoxon two-sided signed-rank test results for encoding models prediction accuracy. WO: word onset, WS: word surprisal, S: spectrogram, E: envelope.

**Table S9:** Univariate contributions to ISC C1 within attentional conditions.

| attentive |  |  |  |  | distracted |  |  |
| --- | --- | --- | --- | --- | --- | --- | --- |
| group1 | group2 | W | p | $p_{adj}^1$ | W | p | $p_{adj}^1$ |
| original data | -WO | 350 | 5.96e-08 | **5.96e-07** | 351 | 2.98e-08 | **1.19e-07** |
| original data | -WS | 346 | 2.98e-07 | **2.68e-06** | 350 | 5.96e-08 | **1.79e-07** |
| original data | -S | 342 | 9.83e-07 | **7.86e-06** | 351 | 2.98e-08 | **1.19e-07** |
| original data | -E | 333 | 7.54e-06 | **5.28e-05** | 351 | 2.98e-08 | **1.19e-07** |
| -WO | -WS | 270 | 1.50e-02 | **3.00e-02** | 324 | 3.76e-05 | **3.76e-05** |
| -WO | -S | 321 | 6.03e-05 | **3.02e-04** | 351 | 2.98e-08 | **1.19e-07** |
| -WO | -E | 297 | 1.00e-03 | **4.00e-03** | 351 | 2.98e-08 | **1.19e-07** |
| -WS | -S | 324 | 3.76e-05 | **2.26e-04** | 351 | 2.98e-08 | **1.19e-07** |
| -WS | -E | 301 | 8.35e-04 | **3.00e-03** | 351 | 2.98e-08 | **1.19e-07** |
| -S | -E | 176 | 1.00e+00 | 1.00e+00 | 345 | 4.17e-07 | **8.34e-07** |

<sup>1</sup> FDR corrected. Bold indicates $p < 0.05$
Wilcoxon two-sided signed-rank test results for ISC of component one after encoding model subtraction. WO: word onset, WS: word surprisal, S: spectrogram, E: envelope. The (-) symbol indicates that the predicted activity by the encoding model was subtracted from the EEG response.

**Table S10:** Univariate contributions to ISC C2 within attentional conditions.

| attentive |  |  |  |  | distracted |  |  |
| --- | --- | --- | --- | --- | --- | --- | --- |
| group1 | group2 | W | p | p<sup>1</sup><sub>adj</sub> | W | p | p<sup>1</sup><sub>adj</sub> |
| original data | -WO | 345 | 4.17e-07 | **3.75e-06** | 313 | 0.000190 | **0.002** |
| original data | -WS | 350 | 5.96e-08 | **5.96e-07** | 274 | 0.011000 | 0.066 |
| original data | -S | 204 | 4.83e-01 | 6.71e-01 | 127 | 0.227000 | 0.908 |
| original data | -E | 261 | 2.90e-02 | 1.46e-01 | 143 | 0.423000 | 0.940 |
| -WO | -WS | 332 | 9.15e-06 | **7.32e-05** | 204 | 0.483000 | 0.940 |
| -WO | -S | 127 | 2.27e-01 | 6.71e-01 | 76 | 0.010000 | 0.066 |
| -WO | -E | 193 | 6.71e-01 | 6.71e-01 | 89 | 0.027000 | 0.136 |
| -WS | -S | 39 | 2.17e-04 | **1.00e-03** | 39 | 0.000217 | **0.002** |
| -WS | -E | 129 | 2.47e-01 | 6.71e-01 | 60 | 0.002000 | **0.019** |
| -S | -E | 317 | 1.09e-04 | **7.63e-04** | 179 | 0.940000 | 0.940 |

<sup>1</sup> FDR corrected. Bold indicates p < 0.05
Wilcoxon two-sided signed-rank test results for ISC of component two after encoding model subtraction. WO: word onset, WS: word surprisal, S: spectrogram, E: envelope. The (-) symbol indicates that the predicted activity by the encoding model was subtracted from the EEG response.

**Fig S6.**
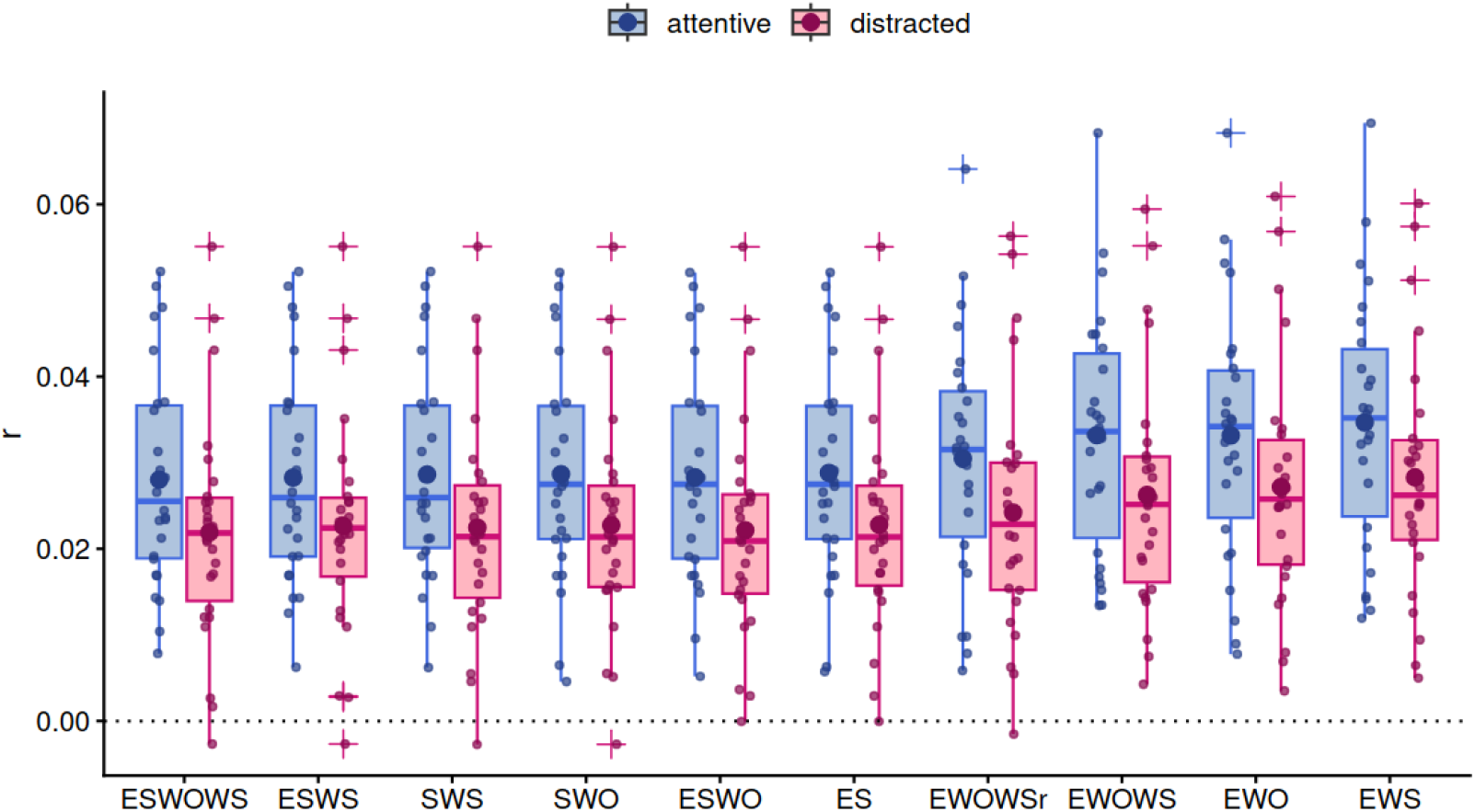
Prediction accuracies for multivariate models are higher in the attentive condition. Multivariate models for combinations of the envelope (E), spectrogram (S), word surprisal (WS), and word onset (WO). Prediction accuracies are higher for the attentive than the distracted condition for all models (p ≤ 0.010). In each boxplot, the horizontal line within the box denotes the median and the dot represents the mean

**Table S11:** Attention effects on the prediction accuracies of the multivariate TRFs.

| x | y | W | p | p<sub>adj</sub><sup>1</sup> |
| --- | --- | --- | --- | --- |
| attentive.ESWOWS | distracted.ESWOWS | 277 | 0.009000 | **0.010** |
| attentive.ESWO | distracted.ESWO | 275 | 0.010000 | **0.010** |
| attentive.ESWS | distracted.ESWS | 278 | 0.008000 | **0.010** |
| attentive.ES | distracted.ES | 282 | 0.006000 | **0.010** |
| attentive.EWS | distracted.EWS | 301 | 0.000835 | **0.007** |
| attentive.EWO | distracted.EWO | 283 | 0.005000 | **0.010** |
| attentive.SWO | distracted.SWO | 282 | 0.006000 | **0.010** |
| attentive.SWS | distracted.SWS | 284 | 0.005000 | **0.010** |
| attentive.EWOWSr | distracted.EWOWSr | 301 | 0.000835 | **0.007** |

<sup>1</sup> FDR corrected. Bold indicates $p < 0.05$
Wilcoxon two-sided signed-rank test of the prediction accuracy of models between attentional conditions

**Fig S7.**
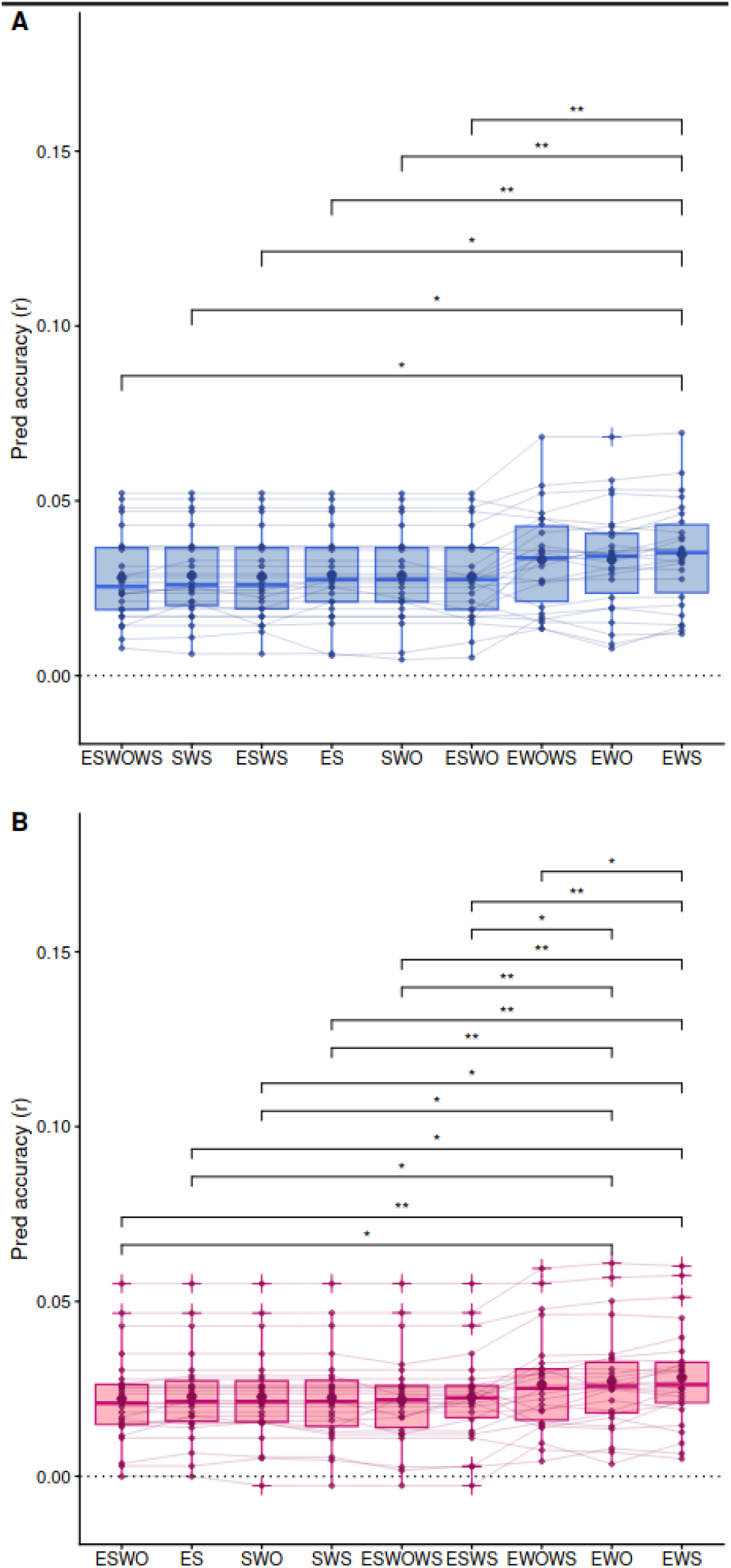
Prediction accuracies for mTRFs models during attentive (**A**, blue) and distracted conditions (**B**, red). Multivariate models for combinations of the envelope (E), spectrogram (S), word surprisal (WS), and word onset (WO). EWO and EWS encoding models show the highest prediction accuracies.

**Table 12:** Multivariate contributions to ISC C1 within attentional conditions.

| attentive |  |  |  |  | distracted |  |  |
| --- | --- | --- | --- | --- | --- | --- | --- |
| x | y | W | p | p<sub>adj</sub><sup>1</sup> | W | p | p<sub>adj</sub><sup>1</sup> |
| original data | -E | 333 | 7.54e-06 | **3.02e-05** | 351 | 2.98e-08 | **1.79e-07** |
| original data | -EWO | 345 | 4.17e-07 | **2.08e-06** | 351 | 2.98e-08 | **1.79e-07** |
| original data | -EWS | 345 | 4.17e-07 | **2.08e-06** | 351 | 2.98e-08 | **1.79e-07** |
| original data | -EWOWS | 345 | 4.17e-07 | **2.08e-06** | 351 | 2.98e-08 | **1.79e-07** |
| -E | -EWO | 351 | 2.98e-08 | **2.68e-07** | 308 | 3.64e-04 | **3.64e-04** |
| -E | -EWS | 351 | 2.98e-08 | **2.68e-07** | 337 | 3.28e-06 | **1.31e-05** |
| -E | -EWOWS | 350 | 5.96e-08 | **4.77e-07** | 344 | 5.66e-07 | **2.83e-06** |
| -EWO | -EWS | 277 | 9.00e-03 | **1.70e-02** | 320 | 7.02e-05 | **1.40e-04** |
| -EWO | -EWOWS | 297 | 1.00e-03 | **4.00e-03** | 351 | 2.98e-08 | **1.79e-07** |
| -EWS | -EWOWS | 233 | 1.50e-01 | 1.50e-01 | 330 | 1.33e-05 | **3.99e-05** |

<sup>1</sup> FDR corrected. Bold indicates $p < 0.05$
Wilcoxon two-sided signed-rank test results for encoding models prediction accuracy. WO: word onset, WS: word surprisal, S: spectrogram, E: envelope.

**Table 13:** Multivariate contributions to ISC C2 within attentional conditions.

| attentive |  |  |  |  | distracted |  |  |
| --- | --- | --- | --- | --- | --- | --- | --- |
| x | y | W | p | padj<sup>1</sup> | W | p | padj<sup>1</sup> |
| original data | -E | 261 | 2.90e-02 | **2.90e-02** | 143 | 4.23e-01 | 9.04e-01 |
| original data | -EWO | 301 | 8.35e-04 | **3.00e-03** | 175 | 1.00e+00 | 1.00e+00 |
| original data | -EWS | 338 | 2.62e-06 | **1.31e-05** | 217 | 3.03e-01 | 9.04e-01 |
| original data | -EWOWS | 336 | 4.08e-06 | **1.63e-05** | 216 | 3.15e-01 | 9.04e-01 |
| -E | -EWO | 345 | 4.17e-07 | **2.50e-06** | 336 | 4.08e-06 | **2.45e-05** |
| -E | -EWS | 351 | 2.98e-08 | **2.38e-07** | 350 | 5.96e-08 | **5.36e-07** |
| -E | -EWOWS | 351 | 2.98e-08 | **2.38e-07** | 350 | 5.96e-08 | **5.36e-07** |
| -EWO | -EWS | 350 | 5.96e-08 | **4.17e-07** | 348 | 1.49e-07 | **1.19e-06** |
| -EWO | -EWOWS | 351 | 2.98e-08 | **2.38e-07** | 346 | 2.98e-07 | **2.09e-06** |
| -EWS | -EWOWS | 266 | 2.00e-02 | **2.90e-02** | 145 | 4.52e-01 | 9.04e-01 |

<sup>1</sup> FDR corrected. Bold indicates $p < 0.05$
Wilcoxon two-sided signed-rank test results for encoding models prediction accuracy. WO: word onset, WS: word surprisal, S: spectrogram, E: envelope.

**Fig S8.**
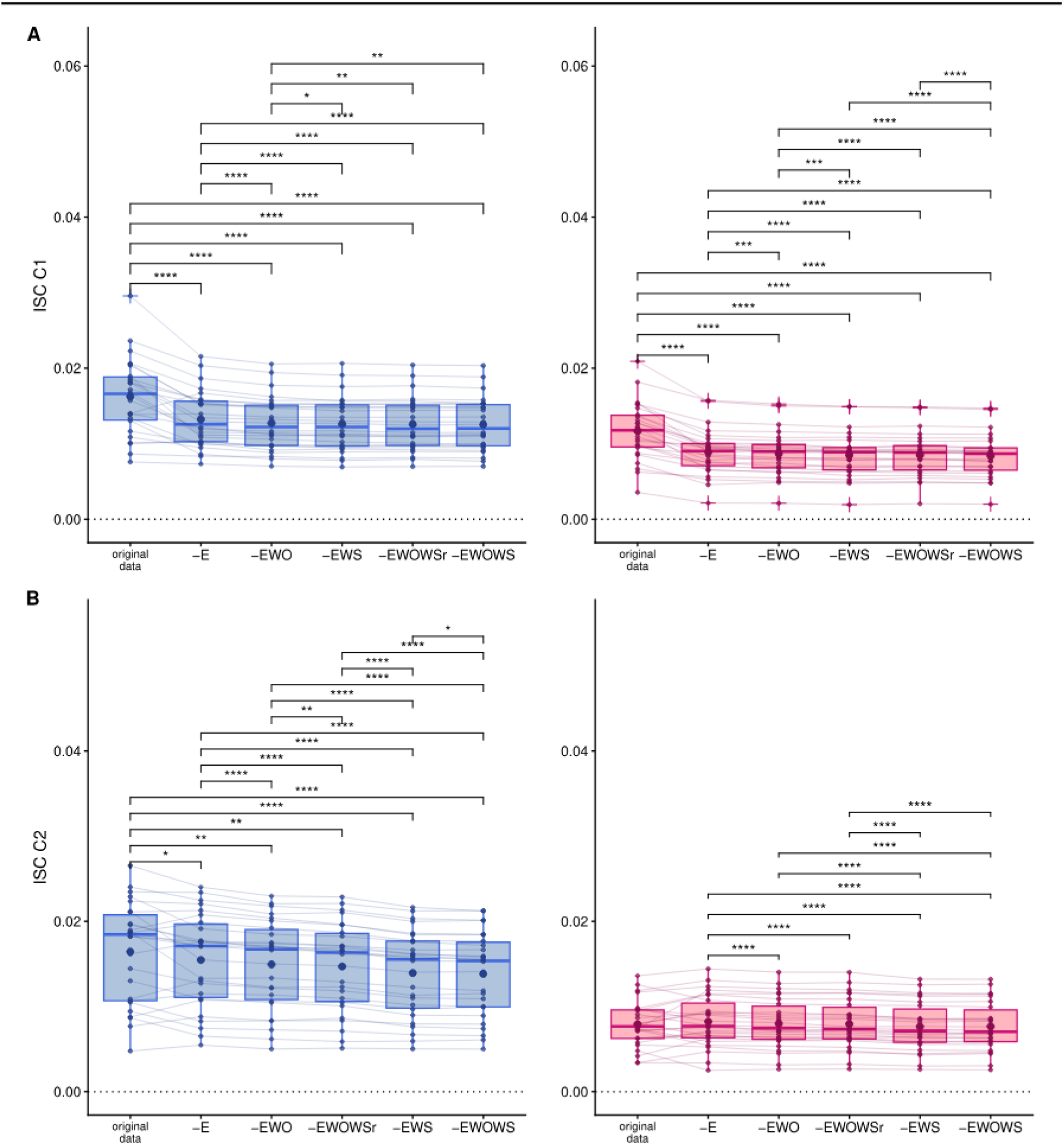
Contributions of mTRFs predictions on the intersubject correlation. A. Component 1. B. Component 2. Multivariate models for combinations of the envelope (E), spectrogram (S), word surprisal (WS), word onset (WO), and randomized word surprisal (WSrand). For the second correlated component, the activity predicted by the EWOWS model contributes more to the ISC than the EWS and EWOWSr models only during the attentive condition.

